# AMR-GNN: A multi-representation graph neural network framework to enable genomic antimicrobial resistance prediction

**DOI:** 10.1101/2025.07.24.666581

**Authors:** Hoai-An Nguyen, Anton Y. Peleg, Jessica A. Wisniewski, Xiaoyu Wang, Zhikang Wang, Luke V. Blakeway, Gnei Z. Badoordeen, Ravali Theegala, Nhu Quynh Doan, Matthew H. Parker, Anna G. Green, Jiangning Song, David L. Dowe, Nenad Macesic

## Abstract

Whole-genome sequencing (WGS) data are an invaluable resource for understanding antimicrobial resistance (AMR) mechanisms. However, WGS data are high-dimensional and the lack of standardized genomic representations is a key barrier to AMR prediction. To fully explore these high-resolution data, we propose AMR-GNN, a graph deep learning-based framework that integrates multiple genomic representations with graph neural networks (GNN) to enable AMR prediction from genomic sequence data. We tested AMR-GNN with *Pseudomonas aeruginosa*, a clinically relevant Gram-negative bacterial pathogen known for its complex AMR mechanisms. We demonstrate that AMR-GNN addresses several key problems in AMR prediction with data-driven machine learning (ML) approaches, including using multiple genomic representations to enhance performance, mitigate the influence of clonal relationships, and identify informative biomarkers to provide explainability and generate novel hypotheses. Follow-up validation on the largest publicly available dataset spanning both Gram-negative and Gram-positive pathogens highlights AMR-GNN’s broad applicability in detecting AMR in diverse and clinically relevant pathogen-drug combinations.

## INTRODUCTION

Antimicrobial resistance (AMR) is a significant global health threat, leading to 1.14 million deaths globally in 2021 [1]. Rapid and accurate detection of AMR through antimicrobial susceptibility testing (AST) is crucial for improving patient outcomes, reducing unnecessary antimicrobial use, and preventing the spread of AMR in hospital settings. Our current methods for identifying AMR are slow, laborious and are often not reported to the clinician for 2-5 days after sample collection [2]. These delays also impact our ability to institute infection prevention measures to stop further AMR transmission.

Whole genome sequencing (WGS) data offer a promising alternative approach to detect AMR. The small size of bacterial genomes makes WGS increasingly accessible to clinical microbiology laboratories. WGS has the potential to streamline processes by replacing multiple individual tests for organism identification and AMR detection [3]. Additionally, the constant generation of public WGS data provides opportunities for surveillance and enhances the robustness of bacterial studies, particularly when statistical analyses often require large sample sizes.

For AMR detection, most pathogens have been analyzed using simple rule-based and/or gene detection approaches [4, 5]. These approaches perform well when AMR is driven by single gene presence or mutations but are less effective in pathogens where AMR results from complex interactions between several factors, or when genetic heterogeneity is present (e.g. recombination or horizontal gene transfer) [3]. They also cannot discover novel genetic variants related to AMR, potentially underutilising the high resolution of WGS data. In recent years, machine learning (ML) has become a promising way to overcome these limitations and has successfully predicted AMR in a wide range of bacterial pathogens [6–9]. ML can account for complex genetic relationships, and once trained, can detect AMR for a broad set of antimicrobials within minutes.

Given the high-dimensional nature of WGS data, bacterial genomes can be represented in multiple ways for input into ML models. However, there is currently no established consensus on the most effective representation [4]. Using several approaches to represent bacterial genomes provides more complete insights into the pathogen. For example, single nucleotide polymorphisms (SNPs) help identify variants relative to a specific reference genome, while pan-genome analysis can uncover variant genes within a population. Despite these potential advantages, the use of multiple genome representations for AMR prediction has remained limited. Recent computational advances (e.g. deep learning) have inspired growing interest in multimodal learning [7, 10, 11], prompting the question of whether this approach can effectively exploit diverse genomic inputs. We therefore aimed to assess the impact of integrating multiple genome representations on AMR prediction, and to leverage modern ML approaches designed for complex, high dimensional data to further enhance prediction performance.

To address these gaps, we identified graph neural networks (GNNs) as a promising solution. Their flexible architecture allows the integration of diverse input types through node and edge features [12]. In contrast to other deep learning architectures, GNNs aggregate information between samples through a similarity network. This characteristic is particularly relevant for AMR prediction as isolates sharing similar genomic features may share similar AST profiles [13]. While GNNs are increasingly applied in biomedical research [12, 14], their use in bacterial genomics for AMR prediction remains limited. We therefore developed AMR-GNN, a new GNN-based framework using multiple genome representations to predict AMR in *P. aeruginosa* as a use case (**Fig. 1a**). This pathogen has a large and diverse genome with high genetic variability and employs a wide range of resistance mechanisms, including efflux pumps, porin loss, and enzymatic inactivation of antibiotics [15, 16]. AMR-GNN outperformed baseline AMR prediction models, including elastic net and convolutional neural network (CNN), across a wide range of antipseudomonal agents. We further evaluated the model on a large, diverse dataset encompassing key Gram-negative and Gram-positive pathogens, confirming AMR-GNN’s effectiveness across multiple species and antimicrobials. Finally, through feature attribution analysis, we highlighted the interpretability of AMR-GNN in identifying AMR biomarkers.

**Fig 1.**
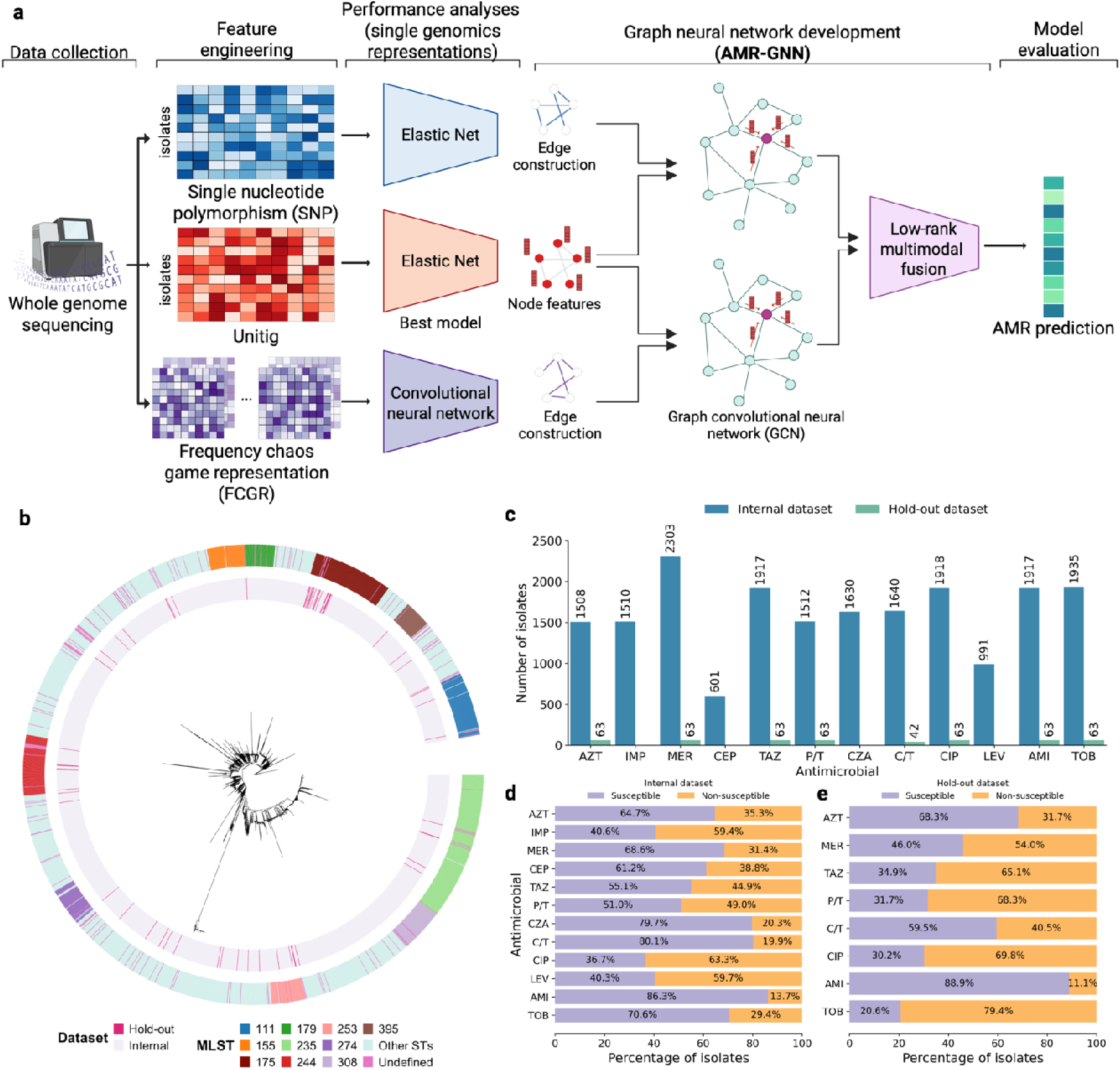
Overview of the AMR-GNN workflow and *P. aeruginosa* dataset used for training/testing. **a.** Schematic representation of AMR-GNN framework. **b.** Phylogenetic tree based on core gene alignment of all *P. aeruginosa* isolates. The inner ring indicates the dataset origin of each isolate. The outer ring represents the multi-locus sequence types (MLSTs). **c.** Number of isolates per antimicrobial in the internal and hold-out dataset. **d, e.** Distribution of susceptible and non-susceptible labels per antimicrobial the internal (d) and hold-out (e) dataset. Abbreviations: AMI: amikacin; AMR: antimicrobial resistance; AZT: aztreonam; C/T: ceftolozane/tazobactam; CEP: cefepime; CIP: ciprofloxacin; CZA: ceftazidime/avibactam; IMP: imipenem; LEV: levofloxacin; MER: meropenem; MLST: multi-locus sequence type; P/T: piperacillin/tazobactam; TAZ: ceftazidime; TOB: tobramycin.

## RESULTS

### Data characteristics

We collected a comprehensive dataset consisting of WGS (Illumina) and AST data from our center and nine publicly available datasets [11, 17–24]. We included datasets with minimum inhibitory concentration (MIC) values obtained from dilution protocols, either broth microdilution (our centre and most public datasets [17–24]) or agar dilution [11]. This resulted in a complete dataset of 2,515 *P. aeruginosa* isolates with diverse population structure (524 unique STs – **Fig. 1b**), with ST235 (10.3%), ST174 (5.5%), ST111 (4.3%) and ST244 (4.0%) most prevalent. AST data included MIC values for 12 common antipseudomonal antimicrobials in binary categories (susceptible/non-susceptible) (**Fig. 1c-e**), per European Committee on Antimicrobial Susceptibility Testing (EUCAST) v15.0 criteria [25]. We used seven datasets (our centre and [11, 17, 19–21, 23, 24]) for training and evaluating the AMR prediction models, with the two smallest datasets [18, 22] kept as hold-out datasets for external validation (see Methods). Non-susceptibility rates varied from 13.7% (amikacin) to 63.3% (ciprofloxacin) in the internal dataset (Fig. 1d), and from 11.1% (amikacin) to 79.4% (tobramycin) in the hold-out datasets (**Fig. 1e**).

### AMR-GNN framework design

AMR-GNN predicts AMR for each isolate by treating each isolate as a node within a graph and performing node classification. Constructing this graph requires two key components: node features represented as feature vectors for each isolate and edge information captured in an adjacency matrix indicating relationships between isolates. We first evaluated AMR prediction using individual genomic representations and selected the best-performing as the node feature input. Following a GNN-based multi-omics integration approach [26], we used the remaining representations to define graph connectivity via adjacency matrices (see **Methods**). Each graph shared the same node features but differed in connectivity, allowing us to learn distinct representations. To integrate information from both graphs, we employed low-rank multimodal fusion to combine their embedding layers into a unified representation used for final AMR prediction [27].

### Models trained with a single genomic representation demonstrated a strong baseline for AMR prediction

We evaluated AMR prediction models trained with different single feature types, including unitigs, SNPs, and frequency chaos game representation (FCGR). For unitigs and SNPs, we applied the elastic net, while FCGR features were used to train a CNN model (see Methods). The unitig and SNP feature sets encompassed 1.5 million and 900,000 features, respectively. For FCGR, we tested k-mer values ranging 5-8 to generate FCGR features and noted that k=7 was optimal, with equivalent performance to k=8 but faster training time (**Supp. Fig. 1**).

Models using unitig features achieved the highest AUROC values across all tested antimicrobials in the internal test sets, showing statistically significant improvements over other feature types for 10/12 antimicrobials, excluding cefepime and ceftolozane/tazobactam (**Fig. 2a, Supp. Table 1**). Unitig-based models exhibited strong predictive performance, particularly for fluoroquinolones (ciprofloxacin AUROC 0.911 [CI: 0.904-0.918]) and aminoglycosides (tobramycin AUROC 0.933 [CI: 0.919-0.947]). Performance for beta-lactams ranged from AUROC 0.878 (CI: 0.847-0.910) for ceftolozane/tazobactam to AUROC 0.636 (CI: 0.603-0.668) for cefepime. When comparing SNP and FCGR representations, FCGR-based models significantly outperformed SNP-based models for ciprofloxacin (0.868 vs 0.733, *P*<0.001), levofloxacin (0.875 vs 0.684, *P*<0.001 and ceftazidime (0.739 vs 0.691, *P*=0.003, with comparable performance across the remaining nine antimicrobials (**Supp. Table 2)**. We then used the salient unitigs selected by the elastic net as node features, while SNP and FCGR data were used to construct the adjacency matrices for AMR-GNN (see Methods).

**Fig 2.**
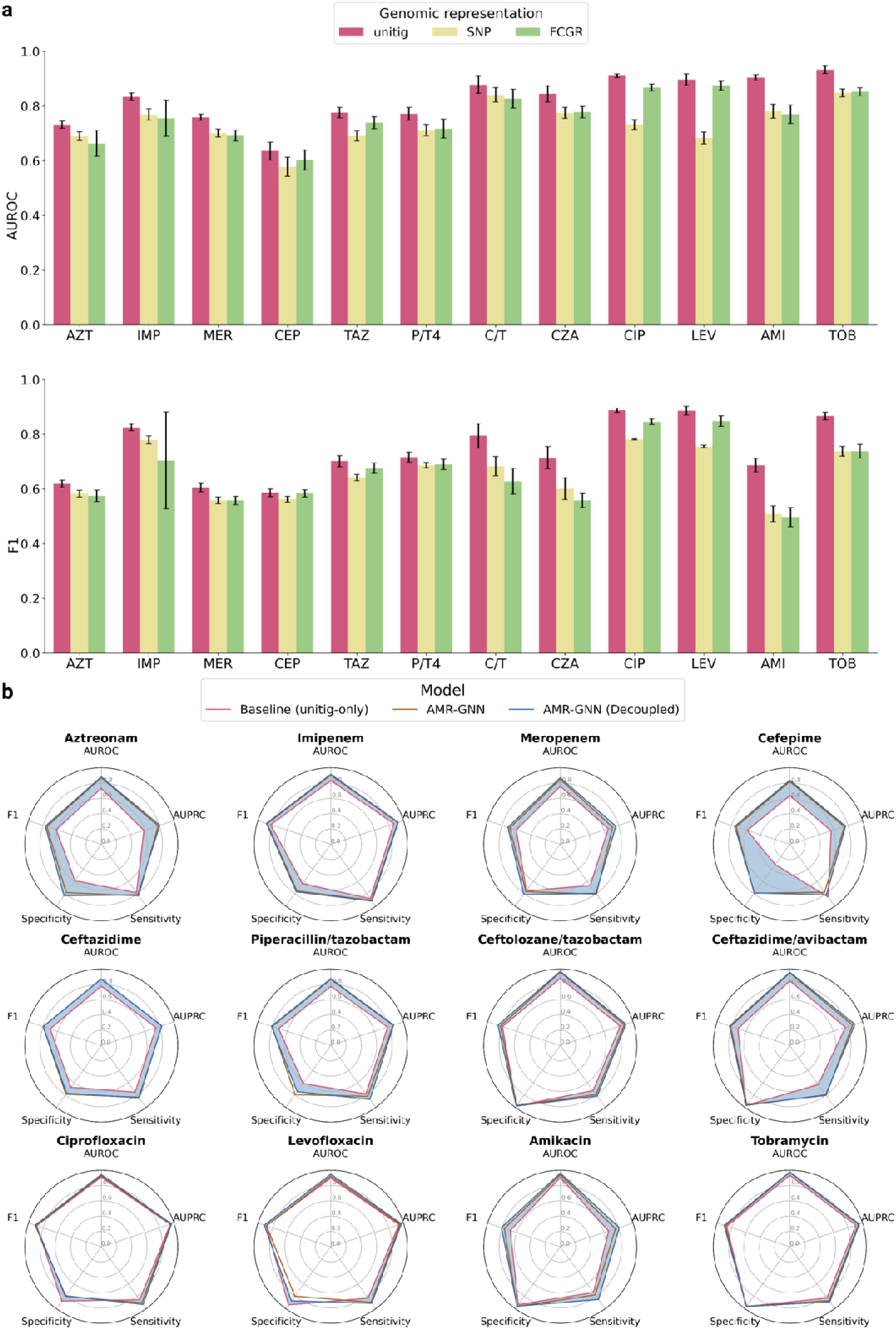
Performance evaluation of individual genomic features and AMR-GNN. **a.** AUROC and F1 performance using individual genomic features (see **Supp. Table 1** for the complete metrics). Bars show the mean performance over 10 random splits with 95% CI error bar. The best-performing single genomic representation (unitig) is selected for comparison with AMR-GNN. **b.** Performance comparison of AMR-GNN (with/without decoupling method) against baseline models (unitig-only models). Color shading denotes the performance difference between the baseline models and the decoupled AMR-GNN, which is our best performing-model. See **Supp. Table 3** for the complete metrics. Abbreviations: AMI: amikacin; AZT: aztreonam; AUROC: area under the receiver operating characteristic curve; AUPRC: area under the precision-recall curve; CEP: cefepime; CIP: ciprofloxacin; C/T: ceftolozane/tazobactam; CZA: ceftazidime/avibactam; FCGR: frequency chaos game representation; IMP: imipenem; LEV: levofloxacin; MER: meropenem; P/T: piperacillin/tazobactam; SNP: single nucleotide polymorphism; TAZ: ceftazidime; TOB: tobramycin.

### AMR-GNN outperformed baseline models

To evaluate the effectiveness of AMR-GNN in improving AMR prediction in *P. aeruginosa*, we used AUROC as the main metric and compared AMR-GNN with unitig-only models as the baseline (**Fig. 2b, Supp. Table 3**). AMR-GNN significantly outperformed unitig-only models in 11/12 tested antimicrobials and performed equivalently for levofloxacin (0.921 vs 0.897, *P*=0.104) (**Supp. Table 4**). Highest AMR-GNN performance was noted for tobramycin (0.971, CI: 0.964-0.978), while lowest was for cefepime (0.819, CI: 0.783-0.855). AMR-GNN made the most substantial improvements for the lowest performing antimicrobials in unitig-only models (cefepime and aztreonam), achieving AUROC increases of 28.8% and 18.9%, respectively.

### Further improving performance by removing bias of bacterial population structure

Bacterial population structure is a major confounder in AMR prediction studies where machine learning models have difficulty differentiating between causal and non-causal variants due to their co-inheritance caused by slow linkage disequilibrium decay [28]. As a result, models may rely on clonal signatures (e.g., high-risk ST clone) to predict AMR instead of uncovering the underlying resistance mechanisms. In the context of GNN, isolates within the same MLST group form dense connections due to their high genetic similarity. This can result in redundant graph information that can degrade performance [29]. Previous work has demonstrated that reducing the influence of redundant neighbors such as edge dropping or node deletion yields more robust results [29, 30]. In this study, we employed a decoupling approach that removes all edges connecting samples from the same MLST group, therefore forcing each node to aggregate information only from neighbors that are not clonally related (see **Methods**). By using this approach, we achieved higher AUROC for all tested antimicrobials compared to the original AMR-GNN, with significant improvements for meropenem, amikacin, and levofloxacin (**Fig. 2b**, **Supp. Table 5**).

### AMR-GNN demonstrates better generalizability on external validation than unitig-only model

After identifying decoupled AMR-GNN as our best-performing approach, we evaluated its performance and benchmarked against untig-only models (our prior baseline comparator) on the hold-out test set for 8 antimicrobials. Compared to the internal test set, AMR-GNN maintained comparable AUROC for ciprofloxacin and ceftazidime, but showed significant performance drops for other antimicrobials (**Supp. Table 6**), suggesting potential overfitting. However, AMR-GNN demonstrated greater robustness to overfitting than unitig-only models, outperforming them in 5/8 antimicrobials tested (**Supp. Table 7**).

### AMR-GNN achieved superior performance compared with state-of-the-art rule-based approaches and publicly available ML models

We benchmarked AMR-GNN against several established approaches (see **Methods**). Rule-based methods are the current biological standard for detecting AMR based on known AMR determinants (including resistance genes and AMR causal variants) [4, 31]. We evaluated a method developed by Cortes-Lara et al. [20], hereafter referred to as the ‘rule-based’ method, using presence/absence of known AMR determinants, and another that combines known AMR determinants and top variants identified by microbial genome-wide association studies and ML-based feature selection to construct an automated scoring system (ARDaP) [32]. In addition, although we previously demonstrated AMR-GNN’s superior performance over elastic net and CNN models, we also compared it with VAMPr, a publicly available AMR prediction tool that uses KEGG (Kyoto Encyclopedia of Genes and Genomes) ortholog variants as features [33]. Since all approaches predict binary outcomes, we evaluated the F1 score (**Fig. 3**).

**Fig 3.**
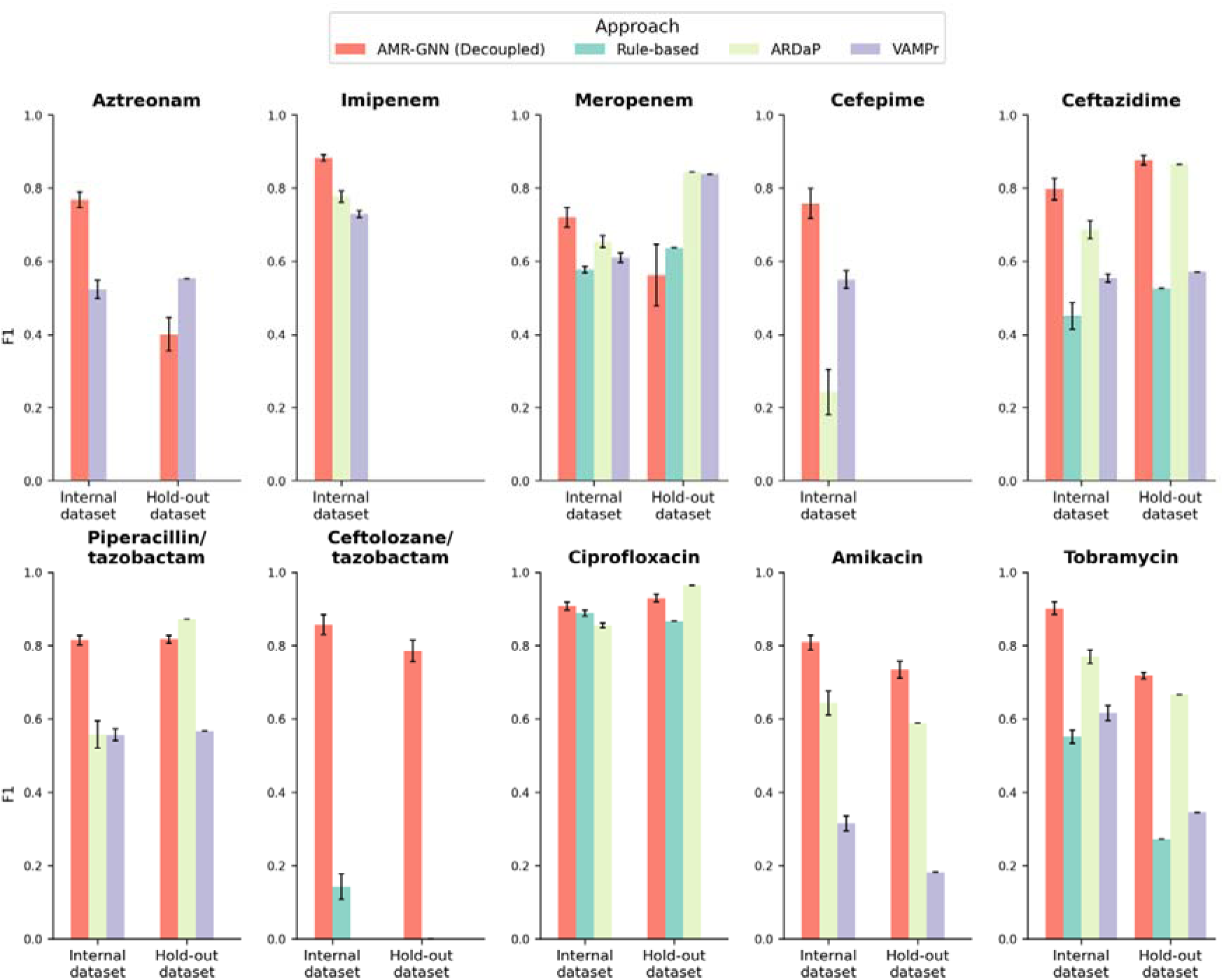
F1-score performance of AMR-GNN compared to rule-based approaches and publicly available AMR prediction models. Performance on both the internal and hold-out test sets is shown. F1-scores are presented as mean with 95% CI error bars.

In the internal dataset evaluation, AMR-GNN demonstrated superior performance with significantly higher F1 scores across all tested antimicrobials compared to benchmark approaches. For the hold-out dataset, AMR-GNN statistically outperformed the rule-based method, ARDaP and VAMPr in 4/5, 3/6 and 4/6 tested antimicrobials, respectively. On the other hand, AMR-GNN underperformed other approaches for meropenem resistance prediction (0.562, CI: 0.478-0.647). We noted meropenem resistance was significantly more prevalent in the hold-out data than in the internal set (54.0% vs 31.4%; P< 0.001, Chi-square test), which likely contributed to this performance drop.

### AMR-GNN demonstrated robust predictive performance across several key bacterial pathogens

To assess the efficacy of AMR beyond *P. aeruginosa*, we evaluated its performance using extensive datasets from two Gram-negative organisms [*Escherichia coli* [n=10,246] and *Klebsiella pneumoniae* [n=7,072]] and two Gram-positive organisms [*Staphylococcus aureus* [n=3,195] and *Enterococcus faecium* [n=2,608]]. The analysis covered a comprehensive panel of antimicrobials specific to each species, sourced from The Pathosystems Resource Integration Center (PATRIC) databases [34, 35] (**Fig. 4a-d**; see **Methods**). AMR-GNN consistently achieved high predictive performance, with mean AUROCs exceeding 0.9 for nearly all tested antimicrobials, except for piperacillin/tazobactam in *E. coli* (0.871, CI: 0.861-0.881) (**Fig. 4e-h**). For each species, highest AUROCs were noted for ceftriaxone in both *E. coli* (0.988; CI: 0.986-0.989) and *K. pneumoniae* (0.985, CI: 0.981-0.990), fusidic acid in *S. aureus* (0.991, CI: 0.988-0.994); and vancomycin in *E. faecium* (0.995, CI: 0.993-0.997). Clinical decision metrics were similarly strong: sensitivity/specificity of 0.938/0.968 for *E. coli*-ceftriaxone, 0.853/0.986 for *K. pneumoniae*-ceftriaxone, 0.990/0.909 for *S. aureus*-fusidic acid, and 0.934/0.992 for *E. faecium*-vancomycin.

**Fig 4.**
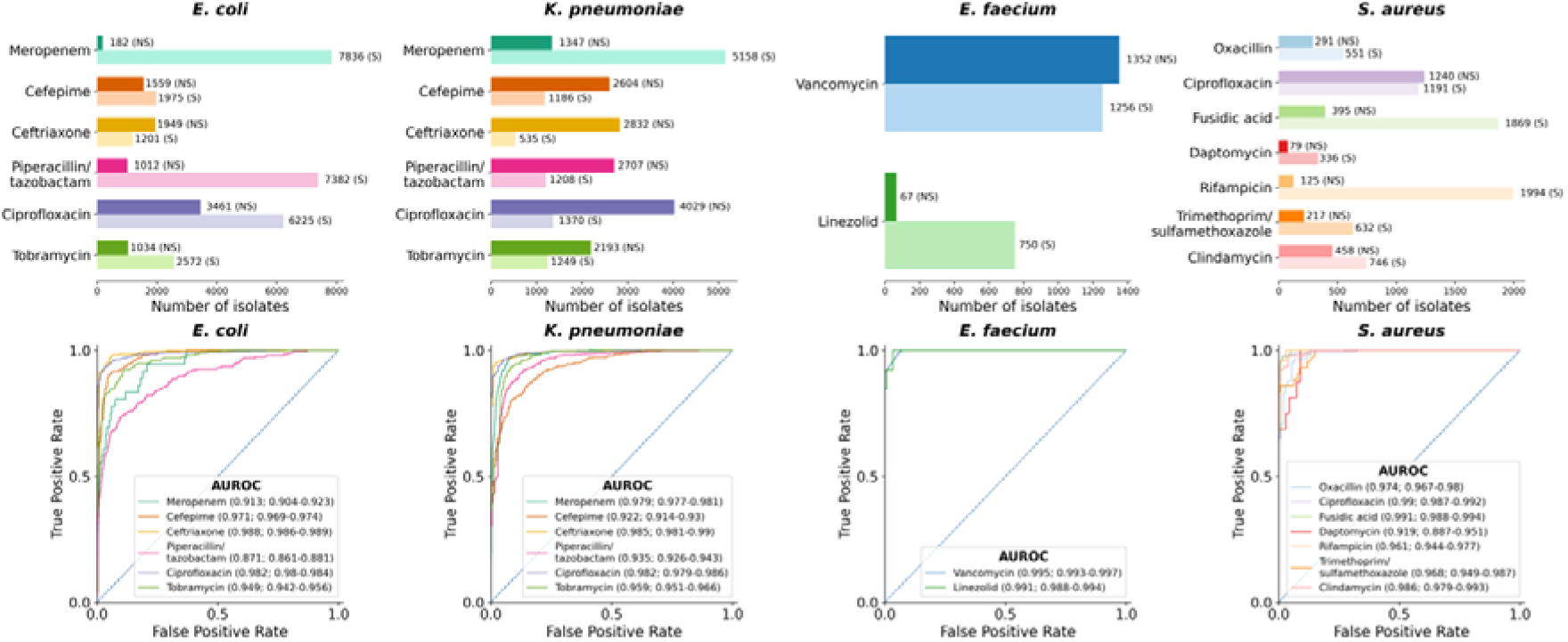
AMR-GNN performance on key bacterial pathogens from the PATRIC database. **a-d.** Phenotypic AST labels (S: susceptible/NS: non-susceptible) of PATRIC dataset across tested antimicrobials for each organism. **e-h.** AUROC performance for each antimicrobial per organism, showing the best curve from 10 random splits.

### Model interpretation on best-performing antimicrobials

We employed integrated gradient (IG) analysis to identify the genetic features contributing most to AMR-GNN performance [36]. We selected the highest-performing antimicrobials from three key antipseudomonal classes: fluoroquinolones (levofloxacin), aminoglycosides (tobramycin), and beta-lactams (ceftolozane/tazobactam) (**Table 1**) [36]. For levofloxacin, the model *a priori* identified known target sites (*gyrA*, *gyrB*, and *parC*) as the most important contributors [37], while PA4179 encodes a probable OpdK porin allowing antimicrobial transport across the bacterial membrane [38, 39]. For tobramycin and ceftolozane/tazobactam, most shortlisted genes had undetermined functions. *fusA1* (tobramycin) has been shown to lead to downstream modification that result in a 4- to 8-fold increase in the MIC for several aminoglycosides, including tobramycin [40]. For ceftolozane/tazobactam, PA4520 is located immediately upstream of *ampE* and *ampD*, which may allow over-expression of AmpC beta-lactamases. We further validated model interpretation results by assessing impact of mutations in shortlisted genes of known functions. Compared to wild-type isolates, MICs were significantly higher in isolates with mutations: *gyrA* (P=2.655 x 10^-134^), *gyrB* (P=6.959 x 10^-5^), *parC* (P=1.053 x 10^-103^) in levofloxacin, and *fusA1* (P= 1.452 x 10^-4^) in tobramycin (Mann-Whitney U-test) (**Supp. Fig. 2a,b**). In terms of specific mutations, we identified 9, 2, and 4 mutations in *gyrA*, *gyrB*, and *parC*, respectively, that were significantly enriched in levofloxacin-resistant isolates (Chi-square test; **Supp. Table 8**). Additionally, 7 mutations in *fusA1* were significantly associated with tobramycin resistance (**Supp. Table 8**).

**Table 1.**
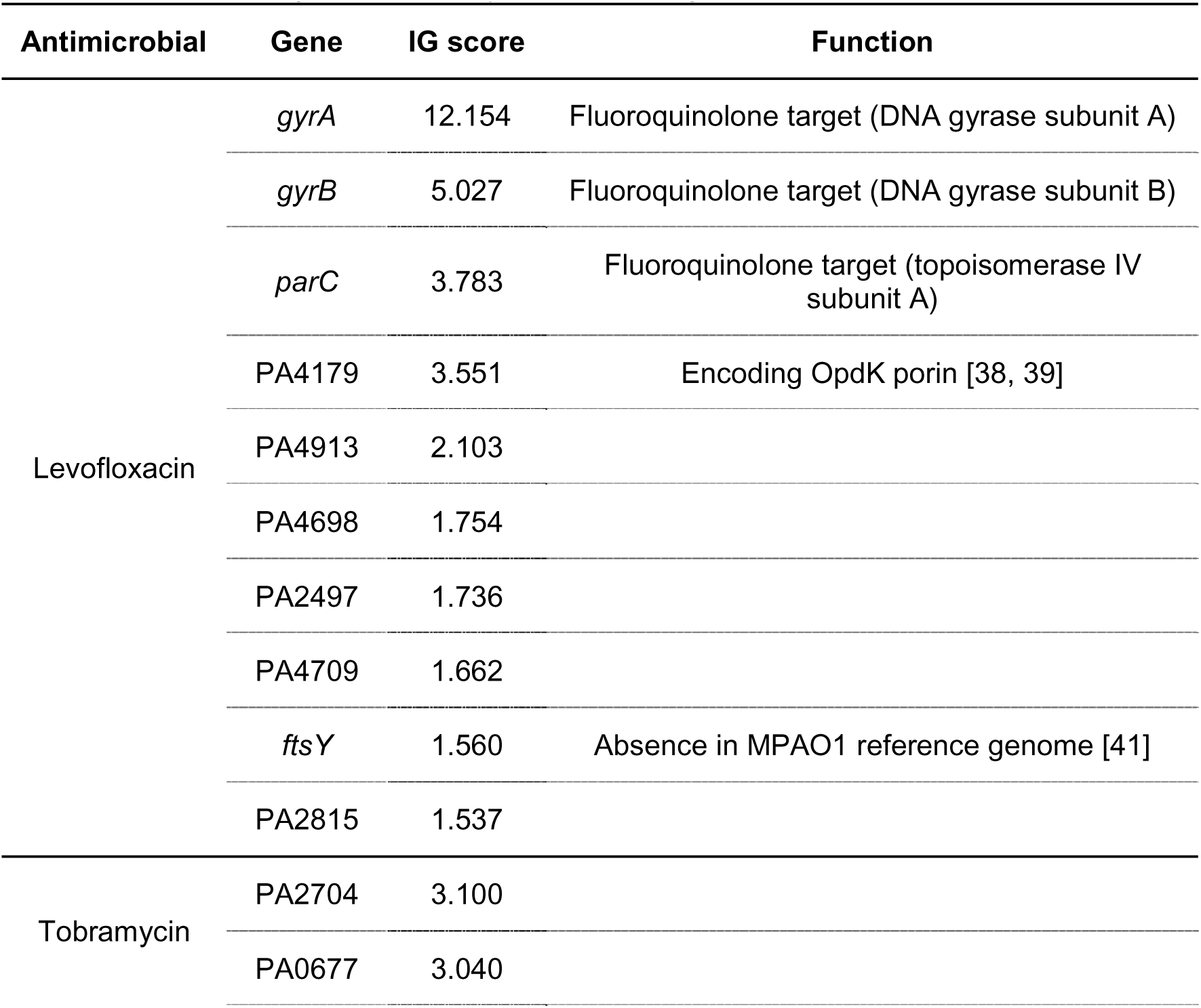
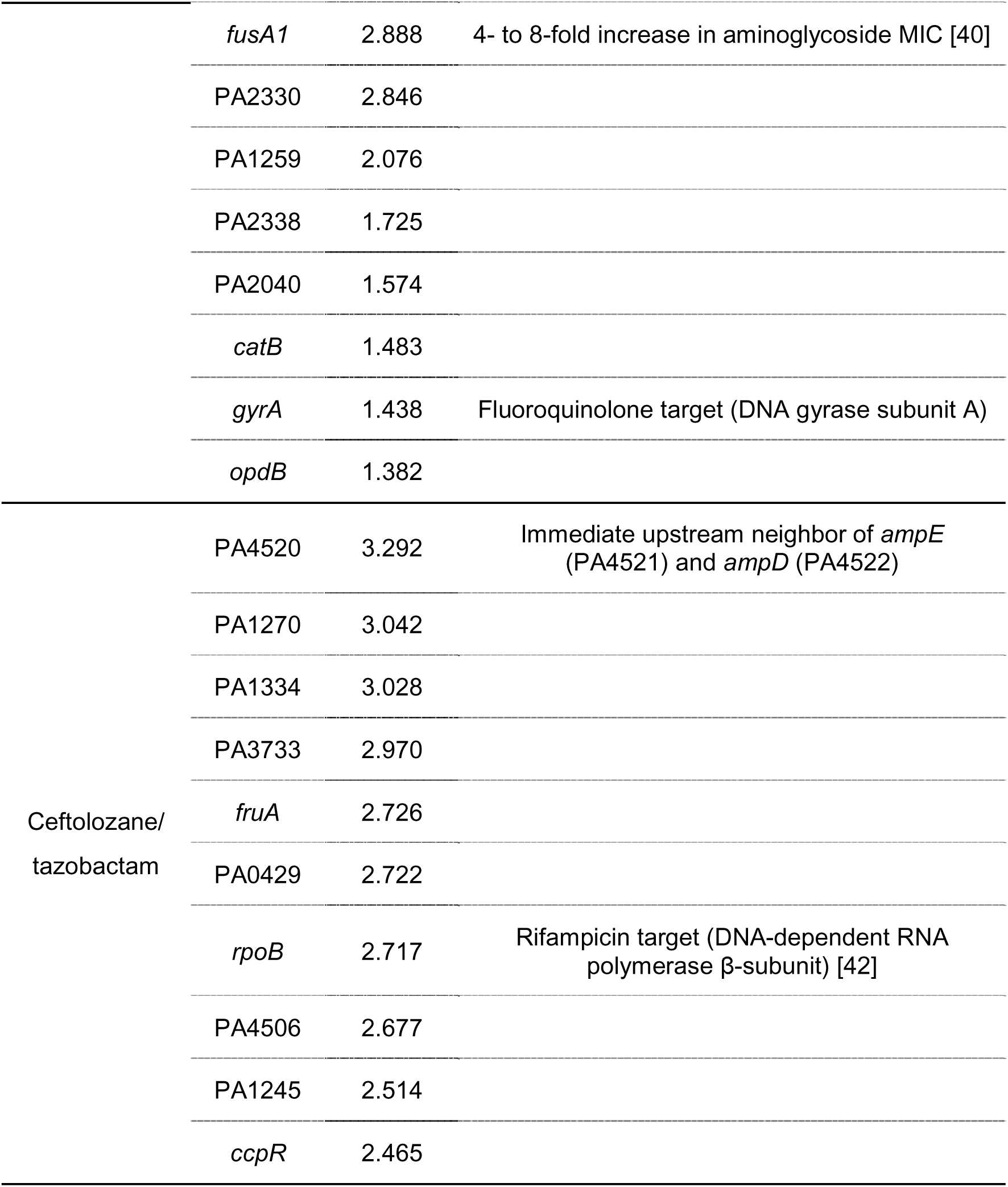
List of the top 10 contributing genes for AMR prediction in levofloxacin, tobramycin, and ceftolozane/tazobactam. For each antimicrobial, genes are displayed in descending order of IG scores.

## DISCUSSION

While ML models leveraging WGS data show promise for AMR prediction, most existing approaches remain limited by their reliance on single genomic representations, failing to fully harness the high-dimensional nature of WGS data. To overcome this limitation, we developed AMR-GNN, a new GNN-based framework that integrates multiple genomic feature sets within a unified deep-learning architecture. This multi-representation strategy achieves substantial performance gains over both conventional ML models and rule-based methods in *P. aeruginosa*, as validated by our benchmarking studies. AMR-GNN leverages two key properties of GNNs: their ability to simultaneously optimise classification outcomes while learning meaningful network structure [14], and their flexibility in processing diverse input data types [12], This enables our model to not only generate informative AMR embeddings but also capture shared genetic content among bacterial isolates, e.g., horizontally transferred AMR genes. Moreover, AMR-GNN also addresses challenges arising from population structure by removing intra-ST edges, enhancing performance robustness. Finally, we showed the species-agnostic transferability of AMR-GNN: once trained on sufficient species-drug pair data, the model accurately predicts resistance patterns across diverse bacterial species and antimicrobial classes.

We demonstrated that AMR-GNN successfully predicts AMR for *P. aeruginosa*, a key nosocomial pathogen with complex mechanisms conferring AMR that have challenged previous AMR prediction efforts. Prior ML efforts were constrained by small cohorts [33], dependence on costly additional data (e.g., transcriptomes) [11], or limited drug coverage [11, 33, 43, 44]. To circumvent these gaps, we curated a large, geographically diverse set of *P. aeruginosa* genomes paired with gold-standard AST results and trained AMR-GNN using multiple, complementary genomic representations. This design consistently outperformed single-feature models and state-of-the-art rule-based approaches across the full spectrum of antipseudomonal agents. Compared with a model that combines WGS and transcriptomics [11], AMR-GNN achieved higher sensitivities for ciprofloxacin (0.931 vs 0.92) and ceftazidime (0.846 vs 0.83), matched performance for tobramycin (0.889 vs 0.89), and was lower for meropenem (0.792 vs 0.91). This is an encouraging trade-off given the expense and limited availability of transcriptomic profiling.

We observed a performance drop in the hold-out dataset, potentially due to differences in susceptibility rates between the internal and hold-out datasets, yet AMR-GNN remained more robust than most benchmarks. It outperformed ARDaP on 3/6 drugs (ceftazidime, amikacin, tobramycin) despite one of the datasets used as hold-out data also having been used for ARDaP training [18, 32]. Relative to the only other study with external validation [45], AMR-GNN proved more robust: its mean AUROC drop across common tested drugs (ciprofloxacin, tobramycin, ceftazidime, meropenem) was 0.076, versus 0.189 in the prior work.

Beyond *P. aeruginosa*, AMR-GNN’s performance across more than 23,000 genomes spanning four clinically important pathogens, tested against a comprehensive panel of species-specific antimicrobials, underscores the framework’s breadth and adaptability. Mean AUROC values surpassed 0.90 for almost every species-drug pair, despite the biological and genomic differences between *E. coli*, *K. pneumoniae*, *S. aureus* and *E. faecium.* This indicates that the graph-based integration of multiple genomic representations, a core design choice of AMR-GNN, successfully captures diverse resistance mechanisms of both Gram-negative bacilli and Gram-positive cocci. Furthermore, we noted the top-scoring combinations (i.e., ceftriaxone in *E. coli,* ciprofloxacin in *K. pneumoniae*, and vancomycin in *E. faecium*) are of significant clinical interest.

Regarding model interpretation, we assessed IG attribution scores to identify salient biomarkers for antimicrobials where AMR-GNN achieved high performance in *P. aeruginosa*. While key AMR drivers were identified in models predicting resistance to levofloxacin and tobramycin, the analysis of ceftolozane/tazobactam resistance revealed several genes of unknown function, emphasizing the complexity of beta-lactam resistance where resistance likely emerges from the interaction of multiple mutations. It might also explain the superiority of AMR-GNN over rule-based approaches in predicting ceftolozane/tazobactam resistance, as the latter relies on known AMR variants. By deriving unitig-based node features without requiring *a priori* AMR knowledge, AMR-GNN eliminates the need for extensive curation of resistance determinants and may facilitate the use of ML approaches for AMR surveillance. Furthermore, the observation of significantly higher MICs in isolates with mutations in the identified AMR determinants further validates the model’s interpretability and biological relevance. These findings not only confirm the effectiveness of our AMR-GNN model in predicting AMR but also underscore its interpretability which is crucial for consideration in clinical applications.

We acknowledge several limitations of our study. First, constructing node features requires a prior feature-selection step, and these chosen features may change as new isolates are added to capture evolving AMR variant profiles. While unitigs for incoming isolates can be derived from the existing database, reconstructing the database entirely may be preferable to avoid falsely missing unitigs that include novel variants. Second, confirming the role of newly identified genes with unknown functions will require laboratory-based validation, which are beyond the scope of this study. Third, despite AMR-GNN’s strong performance, its computational complexity must be weighed against the simplicity of gene-based methods across a wider range of pathogens. We demonstrated this comparison for *P. aeruginosa* but benchmarking of AMR-GNN against other species-specific rule-based approaches was beyond the scope of this study. Finally, a prospective study is needed to assess AMR-GNN’s utility in guiding real-world treatment decisions. Since its performance varies by species–drug combination, we therefore recommend first applying the framework to those combinations with the strongest predictive performance, such as fluoroquinolones and aminoglycosides across multiple species, before broader deployment.

In conclusion, our study demonstrates that the performance of ML-based AMR prediction models can be significantly improved through both the selection of appropriate WGS feature representations and the use of advanced deep learning architectures. Instead of relying on a single feature representation, integrating multiple genomic features offers distinct advantages. AMR-GNN presents an interpretable and flexible framework for such integration, with promising potential for application to multi-omics data as these become increasingly accessible. Although this study focuses solely on WGS data, the modular design of AMR-GNN readily accommodates additional data sources, such as electronic health records for patient network modeling in outbreak detection or transcriptomics data for deeper functional insights.

## METHODS

### Data collection

This study was approved by the Alfred Hospital Ethics Committee (Project No. 185/21). Our dataset comprised short-read (Illumina) genomic data and BMD data of 2,515 *P. aeruginosa* isolates from our center and 9 publicly available datasets (NCBI project accession numbers: PRJEB21341, PRJEB31047, PRJEB37711, PRJEB40140, PRJEB61879, PRJNA264310, PRJNA288601, PRJNA526797, and PRJNA656645) [11, 17–24]. For our in-house collection, BMD was performed using the Sensititre^TM^ Gram negative plate (ThermoFisher, plate code: DKMNG). We converted MIC values to binary label (susceptible/non-susceptible) using the EUCAST v15.0 breakpoint tables [25].

### Training, testing and hold-out dataset generation

First, for external validation, we identified two collections with the smallest sample sizes (42 and 21 isolates) and set them aside as a hold-out test set [18, 22]. The hold-out dataset contained AST results for nine antimicrobials; however, we excluded imipenem from the external validation due to absolute non-susceptible rate (100%). Consequently, we conducted external validation for seven antimicrobials: meropenem, tobramycin, amikacin, ciprofloxacin, ceftolozane/tazobactam, ceftazidime, piperacillin/tazobactam, and aztreonam. We split the remaining data into training and testing sets using an 80/20 stratified split based on AST labels for each antimicrobial. To account for performance variability, this process was repeated 10 times with random seeds. For consistency, the same training/testing splits were used across all experiments, including models based on single genomic representations, AMR-GNN training, and benchmarking with external AMR prediction tools. For AMR-GNN and the CNN model using FCGR inputs, the training set was further divided into training (80%) and validation (20%) subsets. These models were trained on the reduced training set, with hyperparameters tuned based on validation loss. The final models were evaluated on the internal and the hold-out test sets.

### WGS processing

We used Unicycler to assemble short-read WGS data into contigs [46]. The assemblies were then uploaded to Pathogenwatch (https://pathogen.watch) to filter out contaminated isolates, ensuring that only *P. aeruginosa* isolates were included in the analysis, and to obtain MLST results. For unitig input, we generated a unitig database using unitig-caller [47]. To reduce memory requirements associated with large unitig sets, we filtered out unitigs that were either too rare (present in less than 5% of isolates) or too common (present in more than 95%). This filtering reduced the total number of unitigs from 5,590,988 to 1,537,357. Variant calling was performed using Snippy (https://github.com/tseemann/snippy) which generated a SNP profile, including single nucleotide polymorphisms and indels, for each isolate using the PAO1 reference genome (Genbank: NC_002516.1). The individual variant call format (VCF) files were then merged into a single VCF input file using BCFtools for the prediction step with pyseer [48]. FCGR was used to encode the frequency of k-mers into an image-like representation [49]. Following the approach described by Green et al. for *Mycobacterium tuberculosis* [6], we extracted sequences of 57 genes known to be associated with antimicrobial resistance (AMR) in *P. aeruginosa*. The curated gene list can be found in **Supp. Table 9** [15, 20]. The gene sequences were aligned to the PAO1 genome and extracted from the Snippy output. Each gene sequence was then converted into an FCGR matrix, and all matrices were stacked along a new axis. This process resulted in an image-like input with dimensions (C, H, W), where C represents the channel dimension (i.e., the number of AMR genes, n=57), and H=W=k^2^, with k being the size of the k-mer used to extract FCGR features. We tested k values ranging from 5 to 8 and determined that 7-mer was the optimal k-mer setting.

### Machine learning model development for single genomic representation

We trained models to predict AMR using single genomic representations and selected the best-performing representation as our baseline benchmark. Here, we extracted multiple WGS-derived features, including unitig, SNP and FCGR [49]. Such genomic representations have each demonstrated efficacy in AMR prediction and potentially provide different strengths in representing bacterial genomic information [9, 50, 51]. For example, SNPs identify changes in single positions [11] but depend on comparison with a reference genome, while unitigs offer an unbiased, non-reference approach [50, 52]. On the other hand, image-like representations such as FCGR may be more suitable for deep learning models [6, 9, 51]. For unitig and SNP features, we used pyseer’s implementation of the elastic net model with the flags -- wg enet and --alpha 1. For FCGR representation, we used the same CNN architecture as described by Ren et al. [9], consisting 3 convolutional layers and a final classification head. We used the BCELoss function to calculate the loss. We trained our models for 200 epochs with a batch size of 32. We used the Adam optimizer with a multi-step learning rate, starting from 10^-3^ and reducing to 10^-4^ and 10^-5^ after the 50^th^ and 100^th^ epochs, respectively. The best performing model is the one minimizing the validation loss.

### Model evaluation and statistical analysis

We validated our models with 10-fold random splits to ensure robustness. We selected AUROC as our primary metric to assess prediction performance. Additionally, we reported the area under the precision-recall curve (AUPRC), F1-score, sensitivity, specificity, and log-loss. The classification threshold was chosen to maximize the F1-score. All metrics were reported as mean values with 95% confidence intervals (95% CI). Unless otherwise specified, we used the Mann-Whitney U test to compare performance between two different feature types or model architectures. A significance level of *P*=0.05 was applied.

### AMR-GNN development

We employed graph convolutional neural network (GCN) for multi-genomic representation learning [53]. The graph network requires two input types: a feature vector for each node and an adjacency matrix that defines the bi-directional edges connecting pairs of nodes. For each antimicrobial, we constructed the node features using the unitigs selected by elastic net model. Specifically, we retained all unitigs with non-zero beta coefficients. The feature data was represented as a binary feature vector, where a value of 0 indicated the absence of the corresponding unitig in the genome, while a value of 1 denoted its presence.

For edge construction, we first created distance matrices for both SNP and FCGR representations using Hamming code and Euclidean distance, respectively. Next, we arranged the unique distances in ascending order. Considering n as the total number of samples and k‰ as the desired proportion of edges to be retained in the final graph, we calculated the total number of possible undirected edges (|E|_total_) and the number of edges to be kept (|E|_kept_) using the following equations:

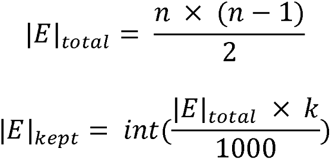

For each distance matrix, we set the distance threshold to determine the adjacency matrix. We incrementally increased the distance threshold, connecting two nodes if their distance was less than the current threshold. At each step, we calculated the total number of edges in the resulting adjacency matrix (|E|_A_). We continued this process until the number of edges exceeded a predefined value |E|_A_ > |E|_kept_. The adjacency matrix obtained from the previous threshold was then selected as the final adjacency matrix. This matrix was converted into the COO (Coordinate list) format, which is compatible with pytorch-geometric’s data object. For the hyperparameter k, we gradually increased the threshold until k=20, as the performance deteriorated significantly beyond that for all antimicrobials. We found that k=2 provided the best results (**Supp. Fig. 3**).

In terms of model architecture, we observed that adjacency matrices derived from different genomic representations varied, particularly at lower retention thresholds (k) (**Supp. Fig. 4**), suggesting that each graph might capture distinct aspects of the data. We therefore developed a dual GCN model that integrates the representation layers from two separate GCN modules into a unified vector input (**Figure 2**). This combined vector serves as the input to predict AMR. Each GCN module learns from the unitig node features and an adjacency matrix, which is inferred from either SNP or FCGR feature. The latent representation *h_i_*, obtained from each GCN module, can be expressed as follows:

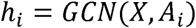

where 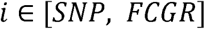, *X* and *A_i_* are the node feature and adjacency matrix, respectively. Specifically, each GCN module consists of two GCN layers, each containing 128 units with rectified linear units (ReLU) activation. We applied layer normalisation and used a dropout rate of 0.5. This generated two 128-dimensional representation vectors prior to the classification head. In literature, various approaches have been explored to combine multiple representations. Among these techniques, tensor presentation has demonstrated superior performance compared to simple fusion methods such as concatenation or pooling. However, such an approach often increases the computational resource exponentially with respect to the number of data modalities. To address this challenge, we adopted the low-rank fusion technique, a tensor representation approach introduced by Liu et al. [27]. This method allows us to efficiently combine the learned representations from two graph networks into the final layer preceding the classification head while maintaining computational efficiency.

### Clonal relationship decoupling process

We first used MLST information to construct a MLST matrix (M) similar to an adjacency matrix, where M_i,j_ =1 if the i^th^ isolate from the row and the j^th^ isolate from the column are in the same MLST group, and M_i,j_ =0 otherwise. We then used the M matrix to update the adjacency matrix (A) such that A_i,j_ =0 if M_i,j_ = 1, resulting in the final decoupled adjacency matrix.

### AMR prediction with rule-based approaches

As no pipeline was provided for the approach by Cortes-Lara et al. [20], we developed our own workflow to classify isolates as susceptible or resistant to five antimicrobials: meropenem, tobramycin, ciprofloxacin, ceftazidime, and ceftolozane/tazobactam. For each antimicrobial, we evaluated the known AMR determinants and assigned scores based on a reference table. Isolates with a cumulative score of 1 or higher were categorized as non-susceptible, while those with scores below 1 were classified as susceptible. We utilized the variant calling results obtained from Snippy to assess chromosomal mutations and queried the genes of interest using the gene annotation results from bakta to determine gene presence or absence [54]. We excluded all natural polymorphism variants, as defined by Cortes-Lara et al., when calculating the scores. Regarding ARDaP [32], we ran the Nextflow pipeline (https://github.com/dsarov/ARDaP) to generate the report and extracted the predicted phenotypes for the tested antimicrobials.

### AMR prediction with VAMPr

We performed command-line VAMPr AMR prediction [33], using the approach described (https://github.com/jiwoongbio/VAMPr). AMR prediction models of amikacin, aztreonam, cefepime, ceftazidime, imipenem, piperacillin/tazobactam, and tobramycin can be found at https://cdc.biohpc.swmed.edu/VAMPr/VAMPr.cgi.

### AMR-GNN validation on PATRIC database

Assemblies of tested isolates were downloaded from the FTP server at ftp://ftp.bvbrc.org [34, 35]. For Gram-negative pathogens (*E. coli* and *K. pneumoniae*), we evaluated susceptibility to cefepime, ceftriaxone, piperacillin/tazobactam, meropenem, and tobramycin. For *E. faecium*, vancomycin and linezolid were tested, whereas for *S. aureus*, we included oxacillin, fusidic acid, ciprofloxacin, clindamycin, daptomycin, rifampicin, and trimethoprim/sulfamethoxazole. To standardize susceptibility labels across datasets, isolates with available MIC values were reclassified according to EUCAST v15 clinical breakpoints [25]. For those having only AST labels, samples labeled as intermediate (“I”) were excluded from analysis as it was unclear which interpretive criteria were used. For each organism-antimicrobial combination, we followed the same approach used previously for *P. aeruginosa*: datasets were split into training and testing subsets, and models were trained across 10 random splits. During feature (unitig) selection for *E. coli*, the training dataset was limited to 3,000 isolates due to memory constraints associated with large variant files. Reference genomes used for variant calling were: *E. coli* str. K-12 substr. MG1655 (Genbank: U00096.3), *K. pneumoniae* subsp. pneumoniae MGH 78578 strain ATCC (Genbank: CP000647.1), *E. faecium* strain SRR24 chromosome (Genbank: CP038996.1), and *S. aureus* subsp. aureus NCTC 8325 (Genbank: CP000253.1). Known resistomes for FCGR feature extraction were derived from the curated Antibiotic Resistance Ontology (ARO) gene list from the Comprehensive Antibiotic Resistance Database (CARD) (**Supp. Table 10-13**) [55].

### Model interpretation

We assessed node feature importance using IG [36]. IG adheres to two key interpretability axioms: (1) sensitivity, ensuring identical attributions for models with identical outputs; and (2) completeness stating that the total attributions equal the output difference between the models using actual input and baseline input (e.g., a zero input). Here, we used Captum’s implementation of IG to compute the attribution scores [56].

To link salient node features (unitigs) to biological meaning, we performed BLAST searches to identify corresponding genes [57]. The top 10 genes were prioritized based on the cumulative IG scores of their mapped unitigs. We then used decision plots derived from SHAP (Shapley Additive Explanation) for interpretation [58].

## Data and source code availability

The in-house sequenced reads have been deposited in the NCBI BioProject ID PRJNA1220180. The Sequence Read Archive (SRA) accession numbers and AST data of all isolates can be found in Supplementary File 1. The source code of this study is available at https://github.com/andyvng/amr-gnn.

## Author contributions

HA.N, A.Y.P. and N.M. conceived the study. J.A.W. designed and supervised sampling and collection of bacterial isolates. J.A.W., L.V.B., G.Z.B, R.T., N.Q.D., and M.H.P. collected the bacterial isolates, performed bacterial characterization, conducted whole genome sequencing and broth microdilution. HA.N. preprocessed data. HA.N., J.S., D.L.D., Z.W and N.M. conceptualized machine learning analyses. HA.N. developed machine learning models and evaluated predictive performances. HA.N. and N.M. analyzed all results. HA.N. and N.M. wrote the manuscript with comments and feedback of all of the co-authors. All authors read and approved the manuscript.

## Acknowledgement

This work was supported by Monash eResearch Center, including the M3 service. This work was also supported by use of the Nectar Research Cloud, a collaborative Australian research platform supported by the National Collaborative Research Infrastructure Strategy (NCRIS)-funded Australian Research Data Commons (ARDC).

