## Supplementary Materials for "AMR-GNN: A multi-representation graph neural network framework to enable genomic antimicrobial resistance prediction"

**Supplementary Figures**

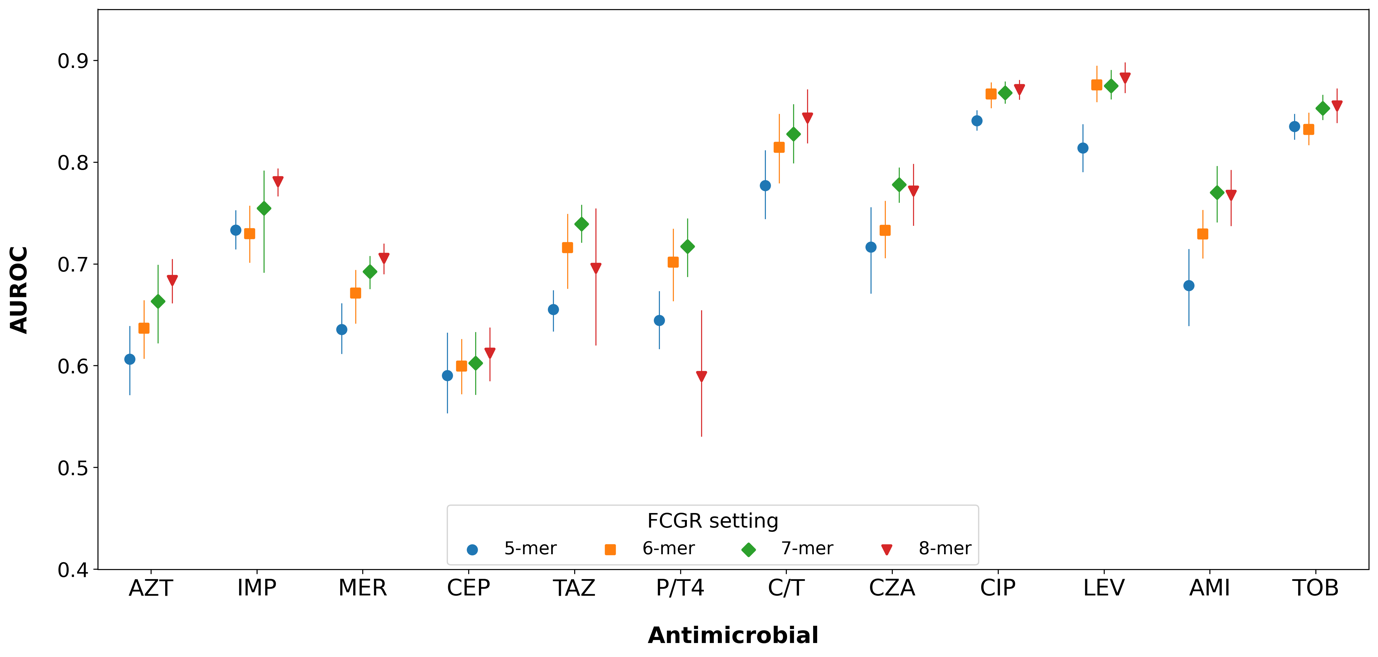

**Supp. Fig. 1.** **AUROC performance per antimicrobial of different k settings to generate FCGR input feature.** Markers and lines indicate mean and 95% confidence interval (CI) of AUROC value, respectively.

Abbreviations: AMI: amikacin; AZT: aztreonam; CEP: cefepime; CIP: ciprofloxacin; C/T: ceftolozane/tazobactam; CZA: ceftazidime/avibactam; FCGR: frequency chaos game representation; IMP: imipenem; LEV: levofloxacin; MER: meropenem; P/T: piperacillin/tazobactam; TAZ: ceftazidime; TOB: tobramycin.

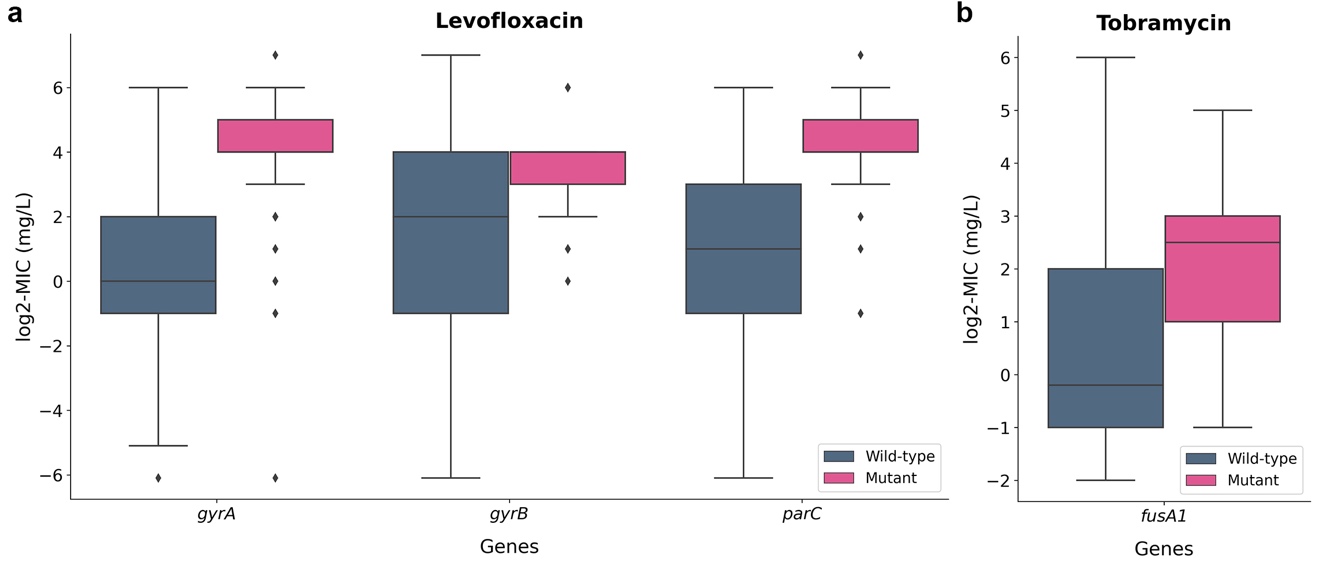

**Supp. Fig 2.** **MIC comparison between wild-type and mutant genes important for AMR prediction.** **a.** Genes *gyrA*, *gyrB*, and *parC* in levofloxacin **b.** Gene *fusA1* in tobramycin.

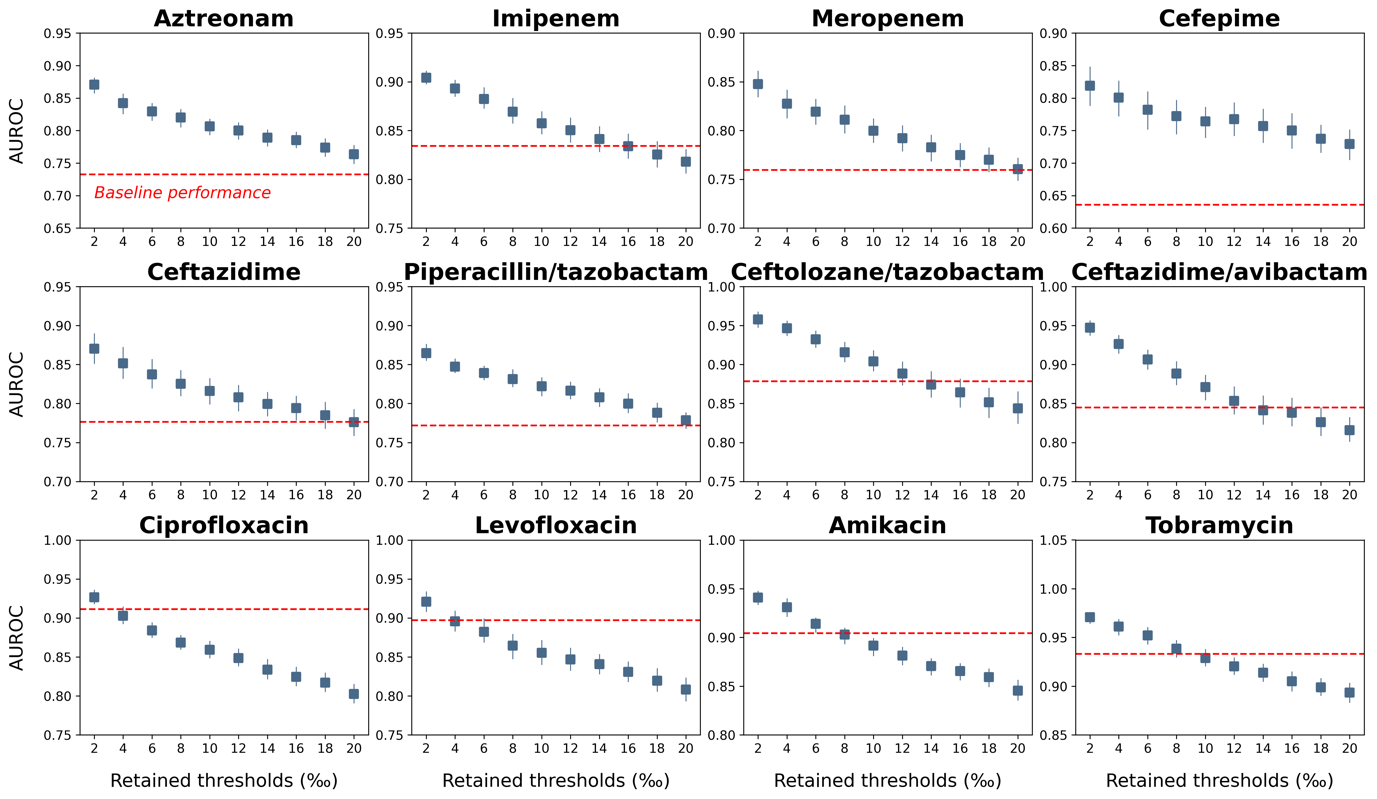

**Supp. Fig. 3.** **AMR performance per antimicrobial at diiferent retained thresholds.**

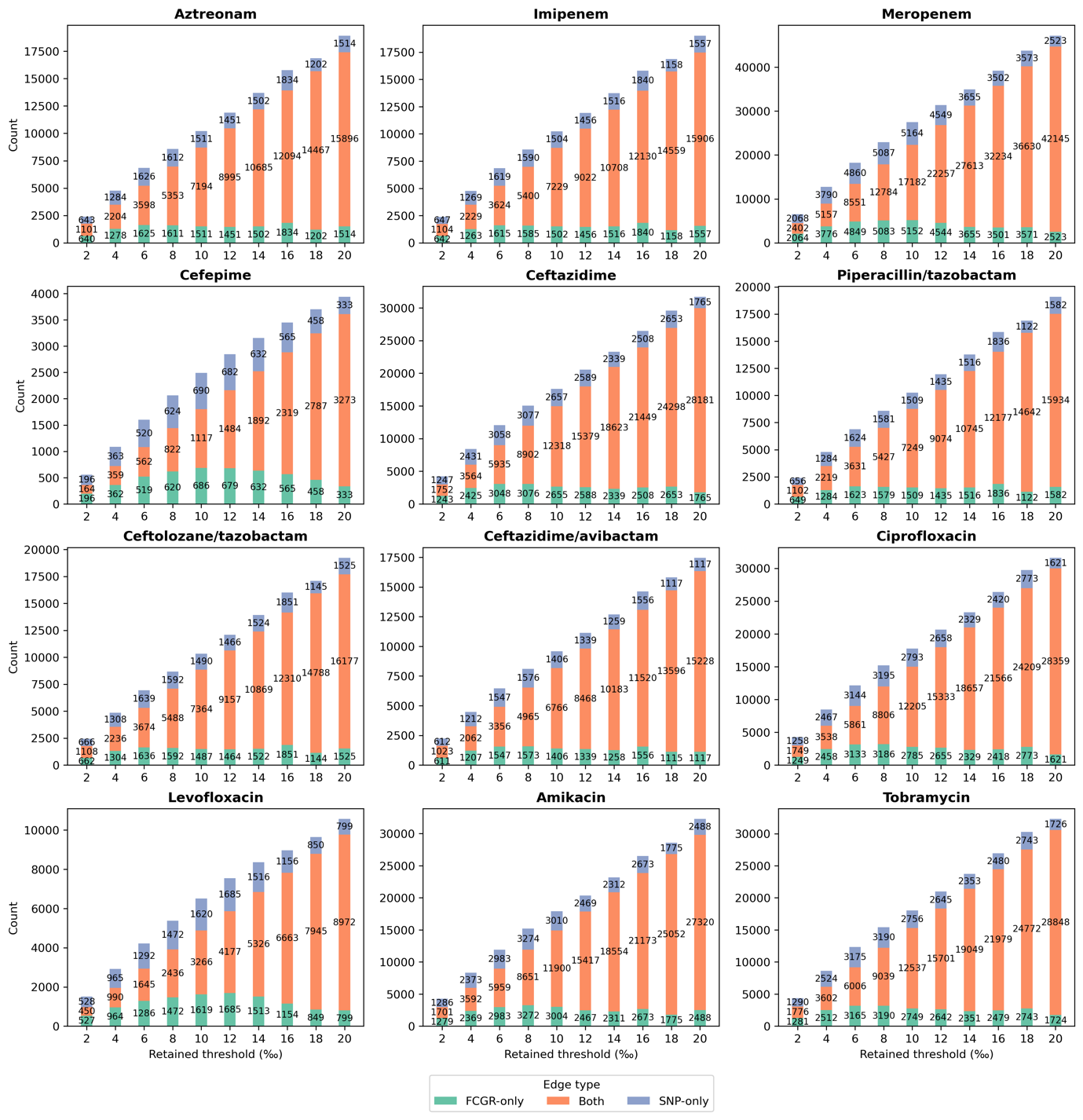

**Supp. Fig. 4. Number of shared and unique edges in adjacency matrices constructed from FCGR- and SNP-based distance matrices at different retained thresholds**

**Supplementary Tables**

**Supp. Table 1. Comprehensive performance metrics for models using individual genomic features.**

Abbreviations: AUROC: area under the receiver operating characteristic curve; AUPRC: area under the precision-recall curve; FCGR: frequency chaos game representation; SNP: short nucleotide polymorphism.

| **Antimicrobial** | **Genomic representation** | **AUROC** | **AUPRC** | **F1** | **Sensitivity** | **Specificity** | **Log-loss** |
| --- | --- | --- | --- | --- | --- | --- | --- |
| Aztreonam | unitig | 0.732 (CI: 0.719-0.745) | 0.587 (CI: 0.557-0.617) | 0.619 (CI: 0.606-0.632) | 0.784 (CI: 0.721-0.848) | 0.587 (CI: 0.505-0.669) | 0.577 (CI: 0.567-0.587) |
|  | SNP | 0.69 (CI: 0.674-0.706) | 0.564 (CI: 0.542-0.585) | 0.583 (CI: 0.571-0.595) | 0.795 (CI: 0.735-0.856) | 0.488 (CI: 0.409-0.567) | 0.599 (CI: 0.592-0.607) |
|  | FCGR | 0.663 (CI: 0.616-0.711) | 0.515 (CI: 0.472-0.557) | 0.575 (CI: 0.554-0.596) | 0.866 (CI: 0.811-0.921) | 0.361 (CI: 0.231-0.491) | 0.618 (CI: 0.602-0.634) |
| Imipenem | unitig | 0.834 (CI: 0.82-0.848) | 0.868 (CI: 0.853-0.882) | 0.825 (CI: 0.813-0.838) | 0.879 (CI: 0.841-0.917) | 0.633 (CI: 0.545-0.722) | 0.5 (CI: 0.485-0.516) |
|  | SNP | 0.769 (CI: 0.748-0.79) | 0.832 (CI: 0.816-0.849) | 0.779 (CI: 0.764-0.794) | 0.896 (CI: 0.855-0.936) | 0.407 (CI: 0.256-0.557) | 0.569 (CI: 0.547-0.59) |
|  | FCGR | 0.755 (CI: 0.689-0.82) | 0.818 (CI: 0.76-0.876) | 0.704 (CI: 0.527-0.882) | 0.762 (CI: 0.564-0.96) | 0.582 (CI: 0.407-0.758) | 0.59 (CI: 0.549-0.632) |
| Meropenem | unitig | 0.759 (CI: 0.748-0.771) | 0.66 (CI: 0.645-0.675) | 0.605 (CI: 0.59-0.62) | 0.666 (CI: 0.608-0.723) | 0.753 (CI: 0.686-0.82) | 0.513 (CI: 0.505-0.522) |
|  | SNP | 0.701 (CI: 0.687-0.716) | 0.576 (CI: 0.557-0.596) | 0.559 (CI: 0.546-0.571) | 0.66 (CI: 0.612-0.708) | 0.675 (CI: 0.605-0.745) | 0.559 (CI: 0.546-0.572) |
|  | FCGR | 0.692 (CI: 0.672-0.712) | 0.504 (CI: 0.476-0.532) | 0.558 (CI: 0.542-0.573) | 0.681 (CI: 0.613-0.749) | 0.649 (CI: 0.56-0.738) | 0.586 (CI: 0.577-0.596) |
| Cefepime | unitig | 0.636 (CI: 0.603-0.668) | 0.569 (CI: 0.534-0.603) | 0.586 (CI: 0.572-0.6) | 0.851 (CI: 0.761-0.941) | 0.326 (CI: 0.167-0.484) | 0.638 (CI: 0.624-0.652) |
|  | SNP | 0.578 (CI: 0.543-0.613) | 0.497 (CI: 0.459-0.534) | 0.563 (CI: 0.552-0.573) | 0.86 (CI: 0.788-0.931) | 0.239 (CI: 0.11-0.369) | 0.66 (CI: 0.644-0.675) |
|  | FCGR | 0.602 (CI: 0.566-0.639) | 0.519 (CI: 0.479-0.559) | 0.584 (CI: 0.571-0.597) | 0.862 (CI: 0.762-0.962) | 0.301 (CI: 0.12-0.483) | 0.657 (CI: 0.645-0.669) |
| Ceftazidime | unitig | 0.776 (CI: 0.757-0.795) | 0.752 (CI: 0.724-0.779) | 0.702 (CI: 0.681-0.722) | 0.754 (CI: 0.697-0.811) | 0.675 (CI: 0.565-0.786) | 0.56 (CI: 0.542-0.579) |
|  | SNP | 0.691 (CI: 0.672-0.709) | 0.68 (CI: 0.655-0.704) | 0.642 (CI: 0.63-0.653) | 0.899 (CI: 0.83-0.969) | 0.267 (CI: 0.142-0.391) | 0.619 (CI: 0.605-0.633) |
|  | FCGR | 0.739 (CI: 0.717-0.761) | 0.71 (CI: 0.682-0.738) | 0.676 (CI: 0.657-0.695) | 0.78 (CI: 0.725-0.836) | 0.567 (CI: 0.451-0.683) | 0.603 (CI: 0.581-0.624) |
| Piperacillin/  tazobactam | unitig | 0.772 (CI: 0.748-0.795) | 0.781 (CI: 0.757-0.806) | 0.716 (CI: 0.698-0.735) | 0.782 (CI: 0.743-0.822) | 0.614 (CI: 0.528-0.699) | 0.565 (CI: 0.546-0.583) |
|  | SNP | 0.712 (CI: 0.691-0.733) | 0.725 (CI: 0.706-0.745) | 0.687 (CI: 0.677-0.697) | 0.883 (CI: 0.82-0.946) | 0.34 (CI: 0.201-0.479) | 0.616 (CI: 0.596-0.636) |
|  | FCGR | 0.717 (CI: 0.683-0.752) | 0.737 (CI: 0.705-0.768) | 0.69 (CI: 0.671-0.709) | 0.899 (CI: 0.842-0.956) | 0.317 (CI: 0.165-0.47) | 0.629 (CI: 0.598-0.66) |
| Ceftolozane/  tazobactam | unitig | 0.878 (CI: 0.847-0.91) | 0.824 (CI: 0.783-0.865) | 0.795 (CI: 0.75-0.839) | 0.728 (CI: 0.675-0.781) | 0.975 (CI: 0.963-0.987) | 0.256 (CI: 0.222-0.289) |
|  | SNP | 0.841 (CI: 0.815-0.868) | 0.737 (CI: 0.701-0.774) | 0.683 (CI: 0.648-0.719) | 0.618 (CI: 0.568-0.669) | 0.953 (CI: 0.939-0.968) | 0.326 (CI: 0.3-0.351) |
|  | FCGR | 0.827 (CI: 0.793-0.862) | 0.63 (CI: 0.581-0.68) | 0.628 (CI: 0.581-0.675) | 0.612 (CI: 0.552-0.673) | 0.915 (CI: 0.883-0.948) | 0.391 (CI: 0.357-0.424) |
| Ceftazidime/  avibactam | unitig | 0.845 (CI: 0.815-0.874) | 0.764 (CI: 0.726-0.802) | 0.714 (CI: 0.674-0.755) | 0.632 (CI: 0.572-0.691) | 0.967 (CI: 0.951-0.982) | 0.312 (CI: 0.283-0.34) |
|  | SNP | 0.775 (CI: 0.755-0.795) | 0.647 (CI: 0.611-0.683) | 0.601 (CI: 0.561-0.641) | 0.523 (CI: 0.474-0.572) | 0.945 (CI: 0.929-0.962) | 0.38 (CI: 0.36-0.4) |
|  | FCGR | 0.778 (CI: 0.757-0.799) | 0.558 (CI: 0.517-0.598) | 0.558 (CI: 0.532-0.583) | 0.598 (CI: 0.536-0.661) | 0.86 (CI: 0.82-0.901) | 0.447 (CI: 0.423-0.472) |
| Ciprofloxacin | unitig | 0.911 (CI: 0.904-0.918) | 0.948 (CI: 0.943-0.954) | 0.888 (CI: 0.88-0.895) | 0.855 (CI: 0.841-0.868) | 0.877 (CI: 0.85-0.905) | 0.349 (CI: 0.334-0.363) |
|  | SNP | 0.733 (CI: 0.715-0.75) | 0.847 (CI: 0.834-0.861) | 0.781 (CI: 0.779-0.783) | 0.982 (CI: 0.97-0.994) | 0.083 (CI: 0.046-0.12) | 0.573 (CI: 0.553-0.592) |
|  | FCGR | 0.868 (CI: 0.855-0.881) | 0.926 (CI: 0.918-0.933) | 0.845 (CI: 0.835-0.856) | 0.82 (CI: 0.782-0.858) | 0.791 (CI: 0.681-0.901) | 0.424 (CI: 0.401-0.447) |
| Levofloxacin | unitig | 0.897 (CI: 0.876-0.918) | 0.936 (CI: 0.916-0.956) | 0.886 (CI: 0.87-0.902) | 0.829 (CI: 0.802-0.856) | 0.939 (CI: 0.912-0.966) | 0.368 (CI: 0.337-0.398) |
|  | SNP | 0.684 (CI: 0.662-0.705) | 0.786 (CI: 0.769-0.802) | 0.755 (CI: 0.751-0.76) | 0.955 (CI: 0.925-0.986) | 0.144 (CI: 0.048-0.239) | 0.624 (CI: 0.614-0.635) |
|  | FCGR | 0.875 (CI: 0.858-0.891) | 0.929 (CI: 0.918-0.94) | 0.848 (CI: 0.829-0.868) | 0.796 (CI: 0.755-0.836) | 0.881 (CI: 0.819-0.944) | 0.417 (CI: 0.383-0.451) |
| Amikacin | unitig | 0.904 (CI: 0.894-0.915) | 0.664 (CI: 0.635-0.692) | 0.687 (CI: 0.662-0.712) | 0.738 (CI: 0.69-0.786) | 0.934 (CI: 0.921-0.948) | 0.245 (CI: 0.232-0.257) |
|  | SNP | 0.781 (CI: 0.756-0.807) | 0.489 (CI: 0.455-0.524) | 0.509 (CI: 0.48-0.538) | 0.475 (CI: 0.404-0.547) | 0.938 (CI: 0.91-0.966) | 0.325 (CI: 0.314-0.335) |
|  | FCGR | 0.77 (CI: 0.736-0.804) | 0.406 (CI: 0.371-0.44) | 0.497 (CI: 0.462-0.532) | 0.566 (CI: 0.511-0.622) | 0.883 (CI: 0.845-0.921) | 0.355 (CI: 0.331-0.379) |
| Tobramycin | unitig | 0.933 (CI: 0.919-0.947) | 0.908 (CI: 0.894-0.922) | 0.867 (CI: 0.852-0.881) | 0.828 (CI: 0.798-0.859) | 0.966 (CI: 0.954-0.977) | 0.248 (CI: 0.231-0.264) |
|  | SNP | 0.848 (CI: 0.833-0.863) | 0.808 (CI: 0.792-0.824) | 0.738 (CI: 0.72-0.756) | 0.671 (CI: 0.635-0.707) | 0.939 (CI: 0.919-0.959) | 0.384 (CI: 0.368-0.4) |
|  | FCGR | 0.853 (CI: 0.838-0.867) | 0.789 (CI: 0.767-0.81) | 0.738 (CI: 0.713-0.763) | 0.689 (CI: 0.648-0.729) | 0.926 (CI: 0.898-0.953) | 0.429 (CI: 0.398-0.461) |

**Supp. Table 2.** **AUROC comparison between models using FCGR and SNP features.** AUROCs are reported as mean with 95% CI.

| **Antimicrobial** | **FCGR** | **SNP** | ***P*** |
| --- | --- | --- | --- |
| Aztreonam | 0.663 (CI: 0.616-0.711) | 0.69 (CI: 0.674-0.706) | 0.623 |
| Imipenem | 0.755 (CI: 0.689-0.82) | 0.769 (CI: 0.748-0.79) | 0.521 |
| Meropenem | 0.692 (CI: 0.672-0.712) | 0.701 (CI: 0.687-0.716) | 0.970 |
| Cefepime | 0.602 (CI: 0.566-0.639) | 0.578 (CI: 0.543-0.613) | 0.307 |
| Ceftazidime | 0.739 (CI: 0.717-0.761) | 0.691 (CI: 0.672-0.709) | 0.004 |
| Piperacillin/tazobactam | 0.717 (CI: 0.683-0.752) | 0.712 (CI: 0.691-0.733) | 0.910 |
| Ceftolozane/tazobactam | 0.827 (CI: 0.793-0.862) | 0.841 (CI: 0.815-0.868) | 0.521 |
| Ceftazidime/avibactam | 0.778 (CI: 0.757-0.799) | 0.775 (CI: 0.755-0.795) | 0.734 |
| Ciprofloxacin | 0.868 (CI: 0.855-0.881) | 0.733 (CI: 0.715-0.75) | <0.001 |
| Levofloxacin | 0.875 (CI: 0.858-0.891) | 0.684 (CI: 0.662-0.705) | <0.001 |
| Amikacin | 0.77 (CI: 0.736-0.804) | 0.781 (CI: 0.756-0.807) | 0.791 |
| Tobramycin | 0.853 (CI: 0.838-0.867) | 0.848 (CI: 0.833-0.863) | 0.623 |

**Supp. Table 3. Comprehensive performance metrics of AMR-GNN (with/without decoupling method) against baseline models (unitig-only models).**

Abbreviations: AUROC: area under the receiver operating characteristic curve; AUPRC: area under the precision-recall curve.

| **Antimicrobial** | **Model** | **AUROC** | **AUPRC** | **F1** | **Sensitivity** | **Specificity** | **Log-loss** |
| --- | --- | --- | --- | --- | --- | --- | --- |
| Aztreonam | Baseline (unitig-only) | 0.732  (CI: 0.719-0.745) | 0.587  (CI: 0.557-0.617) | 0.619  (CI: 0.606-0.632) | 0.784  (CI: 0.721-0.848) | 0.587  (CI: 0.505-0.669) | 0.577  (CI: 0.567-0.587) |
|  | AMR-GNN | 0.87  (CI: 0.856-0.885) | 0.782  (CI: 0.754-0.81) | 0.747  (CI: 0.726-0.768) | 0.831  (CI: 0.784-0.878) | 0.781  (CI: 0.716-0.846) | 0.445  (CI: 0.418-0.473) |
|  | AMR-GNN (Decoupled) | 0.881  (CI: 0.869-0.894) | 0.804  (CI: 0.778-0.829) | 0.768  (CI: 0.746-0.789) | 0.815  (CI: 0.79-0.84) | 0.829  (CI: 0.789-0.869) | 0.45  (CI: 0.414-0.486) |
| Imipenem | Baseline (unitig-only) | 0.834  (CI: 0.82-0.848) | 0.868  (CI: 0.853-0.882) | 0.825  (CI: 0.813-0.838) | 0.879  (CI: 0.841-0.917) | 0.633  (CI: 0.545-0.722) | 0.5  (CI: 0.485-0.516) |
|  | AMR-GNN | 0.904  (CI: 0.896-0.912) | 0.92  (CI: 0.911-0.929) | 0.873  (CI: 0.861-0.885) | 0.906  (CI: 0.885-0.927) | 0.753  (CI: 0.709-0.796) | 0.398  (CI: 0.376-0.42) |
|  | AMR-GNN (Decoupled) | 0.915  (CI: 0.905-0.924) | 0.927  (CI: 0.916-0.938) | 0.883  (CI: 0.874-0.891) | 0.917  (CI: 0.895-0.938) | 0.767  (CI: 0.721-0.813) | 0.395  (CI: 0.363-0.427) |
| Meropenem | Baseline (unitig-only) | 0.759  (CI: 0.748-0.771) | 0.66  (CI: 0.645-0.675) | 0.605  (CI: 0.59-0.62) | 0.666  (CI: 0.608-0.723) | 0.753  (CI: 0.686-0.82) | 0.513  (CI: 0.505-0.522) |
|  | AMR-GNN | 0.848  (CI: 0.831-0.864) | 0.719  (CI: 0.691-0.746) | 0.694  (CI: 0.676-0.713) | 0.804  (CI: 0.761-0.847) | 0.764  (CI: 0.717-0.811) | 0.451  (CI: 0.43-0.472) |
|  | AMR-GNN (Decoupled) | 0.869  (CI: 0.851-0.888) | 0.764  (CI: 0.732-0.797) | 0.72  (CI: 0.694-0.746) | 0.792  (CI: 0.746-0.838) | 0.811  (CI: 0.759-0.863) | 0.436  (CI: 0.411-0.462) |
| Cefepime | Baseline (unitig-only) | 0.636  (CI: 0.603-0.668) | 0.569  (CI: 0.534-0.603) | 0.586  (CI: 0.572-0.6) | 0.851  (CI: 0.761-0.941) | 0.326  (CI: 0.167-0.484) | 0.638  (CI: 0.624-0.652) |
|  | AMR-GNN | 0.819  (CI: 0.783-0.855) | 0.749  (CI: 0.699-0.798) | 0.737  (CI: 0.697-0.777) | 0.774  (CI: 0.728-0.821) | 0.791  (CI: 0.734-0.847) | 0.538  (CI: 0.483-0.592) |
|  | AMR-GNN (Decoupled) | 0.837  (CI: 0.797-0.877) | 0.766  (CI: 0.71-0.822) | 0.758  (CI: 0.717-0.798) | 0.815  (CI: 0.765-0.865) | 0.785  (CI: 0.737-0.833) | 0.516  (CI: 0.466-0.566) |
| Ceftazidime | Baseline (unitig-only) | 0.776  (CI: 0.757-0.795) | 0.752  (CI: 0.724-0.779) | 0.702  (CI: 0.681-0.722) | 0.754  (CI: 0.697-0.811) | 0.675  (CI: 0.565-0.786) | 0.56  (CI: 0.542-0.579) |
|  | AMR-GNN | 0.87  (CI: 0.847-0.894) | 0.833  (CI: 0.801-0.864) | 0.799  (CI: 0.775-0.823) | 0.837  (CI: 0.808-0.867) | 0.787  (CI: 0.717-0.856) | 0.496  (CI: 0.421-0.571) |
|  | AMR-GNN (Decoupled) | 0.869  (CI: 0.84-0.899) | 0.831  (CI: 0.792-0.869) | 0.797  (CI: 0.768-0.826) | 0.846  (CI: 0.813-0.878) | 0.772  (CI: 0.705-0.838) | 0.512  (CI: 0.426-0.597) |
| Piperacillin/  tazobactam | Baseline (unitig-only) | 0.772  (CI: 0.748-0.795) | 0.781  (CI: 0.757-0.806) | 0.716  (CI: 0.698-0.735) | 0.782  (CI: 0.743-0.822) | 0.614  (CI: 0.528-0.699) | 0.565  (CI: 0.546-0.583) |
|  | AMR-GNN | 0.865  (CI: 0.852-0.877) | 0.856  (CI: 0.838-0.874) | 0.806  (CI: 0.795-0.817) | 0.821  (CI: 0.786-0.856) | 0.793  (CI: 0.74-0.845) | 0.498  (CI: 0.457-0.538) |
|  | AMR-GNN (Decoupled) | 0.873  (CI: 0.858-0.888) | 0.864  (CI: 0.844-0.885) | 0.815  (CI: 0.802-0.828) | 0.87  (CI: 0.836-0.904) | 0.746  (CI: 0.703-0.79) | 0.505  (CI: 0.456-0.555) |
| Ceftolozane/  tazobactam | Baseline (unitig-only) | 0.878  (CI: 0.847-0.91) | 0.824  (CI: 0.783-0.865) | 0.795  (CI: 0.75-0.839) | 0.728  (CI: 0.675-0.781) | 0.975  (CI: 0.963-0.987) | 0.256  (CI: 0.222-0.289) |
|  | AMR-GNN | 0.958  (CI: 0.946-0.97) | 0.88  (CI: 0.852-0.909) | 0.822  (CI: 0.795-0.85) | 0.794  (CI: 0.761-0.827) | 0.966  (CI: 0.955-0.977) | 0.242  (CI: 0.191-0.293) |
|  | AMR-GNN (Decoupled) | 0.968  (CI: 0.958-0.979) | 0.902  (CI: 0.874-0.93) | 0.857  (CI: 0.83-0.885) | 0.818  (CI: 0.783-0.854) | 0.978  (CI: 0.968-0.987) | 0.249  (CI: 0.181-0.318) |
| Ceftazidime/  avibactam | Baseline (unitig-only) | 0.845  (CI: 0.815-0.874) | 0.764  (CI: 0.726-0.802) | 0.714  (CI: 0.674-0.755) | 0.632  (CI: 0.572-0.691) | 0.967  (CI: 0.951-0.982) | 0.312  (CI: 0.283-0.34) |
|  | AMR-GNN | 0.947  (CI: 0.935-0.959) | 0.862  (CI: 0.818-0.906) | 0.813  (CI: 0.782-0.845) | 0.806  (CI: 0.769-0.844) | 0.955  (CI: 0.94-0.97) | 0.26  (CI: 0.216-0.305) |
|  | AMR-GNN (Decoupled) | 0.957  (CI: 0.947-0.966) | 0.894  (CI: 0.87-0.918) | 0.827  (CI: 0.796-0.858) | 0.797  (CI: 0.736-0.858) | 0.968  (CI: 0.955-0.981) | 0.271  (CI: 0.217-0.326) |
| Ciprofloxacin | Baseline (unitig-only) | 0.911  (CI: 0.904-0.918) | 0.948  (CI: 0.943-0.954) | 0.888  (CI: 0.88-0.895) | 0.855  (CI: 0.841-0.868) | 0.877  (CI: 0.85-0.905) | 0.349  (CI: 0.334-0.363) |
|  | AMR-GNN | 0.927  (CI: 0.915-0.938) | 0.958  (CI: 0.95-0.965) | 0.896  (CI: 0.884-0.907) | 0.905  (CI: 0.881-0.929) | 0.8  (CI: 0.746-0.854) | 0.371  (CI: 0.335-0.407) |
|  | AMR-GNN (Decoupled) | 0.943  (CI: 0.933-0.953) | 0.966  (CI: 0.96-0.973) | 0.908  (CI: 0.898-0.917) | 0.931  (CI: 0.913-0.949) | 0.792  (CI: 0.754-0.83) | 0.353  (CI: 0.31-0.396) |
| Levofloxacin | Baseline (unitig-only) | 0.897  (CI: 0.876-0.918) | 0.936  (CI: 0.916-0.956) | 0.886  (CI: 0.87-0.902) | 0.829  (CI: 0.802-0.856) | 0.939  (CI: 0.912-0.966) | 0.368  (CI: 0.337-0.398) |
|  | AMR-GNN | 0.921  (CI: 0.905-0.937) | 0.952  (CI: 0.941-0.962) | 0.883  (CI: 0.866-0.899) | 0.897  (CI: 0.872-0.922) | 0.798  (CI: 0.726-0.869) | 0.365  (CI: 0.318-0.412) |
|  | AMR-GNN (Decoupled) | 0.948  (CI: 0.93-0.966) | 0.97  (CI: 0.96-0.98) | 0.913  (CI: 0.891-0.935) | 0.911  (CI: 0.878-0.944) | 0.874  (CI: 0.835-0.912) | 0.327  (CI: 0.245-0.41) |
| Amikacin | Baseline (unitig-only) | 0.904  (CI: 0.894-0.915) | 0.664  (CI: 0.635-0.692) | 0.687  (CI: 0.662-0.712) | 0.738  (CI: 0.69-0.786) | 0.934  (CI: 0.921-0.948) | 0.245  (CI: 0.232-0.257) |
|  | AMR-GNN | 0.941  (CI: 0.932-0.949) | 0.758  (CI: 0.725-0.79) | 0.769  (CI: 0.743-0.796) | 0.777  (CI: 0.725-0.83) | 0.962  (CI: 0.957-0.966) | 0.213  (CI: 0.191-0.234) |
|  | AMR-GNN (Decoupled) | 0.962  (CI: 0.955-0.97) | 0.81  (CI: 0.774-0.847) | 0.809  (CI: 0.789-0.829) | 0.851  (CI: 0.801-0.9) | 0.96  (CI: 0.948-0.971) | 0.19  (CI: 0.165-0.215) |
| Tobramycin | Baseline (unitig-only) | 0.933  (CI: 0.919-0.947) | 0.908  (CI: 0.894-0.922) | 0.867  (CI: 0.852-0.881) | 0.828  (CI: 0.798-0.859) | 0.966  (CI: 0.954-0.977) | 0.248  (CI: 0.231-0.264) |
|  | AMR-GNN | 0.971  (CI: 0.964-0.978) | 0.944  (CI: 0.934-0.955) | 0.887  (CI: 0.877-0.898) | 0.872  (CI: 0.854-0.89) | 0.961  (CI: 0.953-0.969) | 0.244  (CI: 0.202-0.285) |
|  | AMR-GNN (Decoupled) | 0.977  (CI: 0.972-0.982) | 0.955  (CI: 0.946-0.964) | 0.902  (CI: 0.885-0.919) | 0.889  (CI: 0.866-0.913) | 0.965  (CI: 0.957-0.974) | 0.236  (CI: 0.191-0.281) |

**Supp. Table 4. AUROC comparison between unitig-only model and AMR-GNN.** AUROCs are reported as mean with 95% CI.

| **Antimicrobial** | **Unitig** | **AMR-GNN** | ***P*** |
| --- | --- | --- | --- |
| Aztreonam | **0.732 (CI: 0.719-0.745)** | **0.87 (CI: 0.856-0.885)** | **<0.001** |
| Imipenem | **0.834 (CI: 0.82-0.848)** | **0.904 (CI: 0.896-0.912)** | **<0.001** |
| Meropenem | **0.759 (CI: 0.748-0.771)** | **0.848 (CI: 0.831-0.864)** | **<0.001** |
| Cefepime | **0.636 (CI: 0.603-0.668)** | **0.819 (CI: 0.783-0.855)** | **<0.001** |
| Ceftazidime | **0.776 (CI: 0.757-0.795)** | **0.87 (CI: 0.847-0.894)** | **<0.001** |
| Piperacillin/tazobactam | **0.772 (CI: 0.748-0.795)** | **0.865 (CI: 0.852-0.877)** | **<0.001** |
| Ceftolozane/tazobactam | **0.878 (CI: 0.847-0.91)** | **0.958 (CI: 0.946-0.97)** | **<0.001** |
| Ceftazidime/avibactam | **0.845 (CI: 0.815-0.874)** | **0.947 (CI: 0.935-0.959)** | **<0.001** |
| Ciprofloxacin | **0.911 (CI: 0.904-0.918)** | **0.927 (CI: 0.915-0.938)** | **0.017** |
| Levofloxacin | 0.897 (CI: 0.876-0.918) | 0.921 (CI: 0.905-0.937) | 0.104 |
| Amikacin | **0.904 (CI: 0.894-0.915)** | **0.941 (CI: 0.932-0.949)** | **<0.001** |
| Tobramycin | **0.933 (CI: 0.919-0.947)** | **0.971 (CI: 0.964-0.978)** | **<0.001** |

**Supp. Table 5. AUROC comparison between AMR-GNN with and without decoupling approach.** AUROCs are reported as mean with 95% CI.

| **Antimicrobial** | **AMR-GNN (Decoupled)** | **AMR-GNN (Original)** | ***P*** |
| --- | --- | --- | --- |
| Aztreonam | 0.881 (CI: 0.869-0.894) | 0.87 (CI: 0.856-0.885) | 0.212 |
| Imipenem | 0.915 (CI: 0.905-0.924) | 0.904 (CI: 0.896-0.912) | 0.089 |
| Meropenem | **0.869 (CI: 0.851-0.888)** | **0.848 (CI: 0.831-0.864)** | **0.045** |
| Cefepime | 0.837 (CI: 0.797-0.877) | 0.819 (CI: 0.783-0.855) | 0.364 |
| Ceftazidime | 0.869 (CI: 0.84-0.899) | 0.87 (CI: 0.847-0.894) | 0.970 |
| Piperacillin/tazobactam | 0.873 (CI: 0.858-0.888) | 0.865 (CI: 0.852-0.877) | 0.162 |
| Ceftolozane/tazobactam | 0.968 (CI: 0.958-0.979) | 0.958 (CI: 0.946-0.97) | 0.212 |
| Ceftazidime/avibactam | 0.957 (CI: 0.947-0.966) | 0.947 (CI: 0.935-0.959) | 0.186 |
| Ciprofloxacin | 0.943 (CI: 0.933-0.953) | 0.927 (CI: 0.915-0.938) | 0.076 |
| Levofloxacin | **0.948 (CI: 0.93-0.966)** | **0.921 (CI: 0.905-0.937)** | **0.014** |
| Amikacin | **0.962 (CI: 0.955-0.97)** | **0.941 (CI: 0.932-0.949)** | **0.003** |
| Tobramycin | **0.977 (CI: 0.972-0.982)** | 0.971 (CI: 0.964-0.978) | 0.140 |

**Supp. Table 6. AUROC comparison of decoupled AMR-GNN for internal and hold-out test set.** AUROCs are reported as mean with 95% CI.

| **Antimicrobial** | **Internal test set** | **Hold-out test set** | ***P*** |
| --- | --- | --- | --- |
| Aztreonam | **0.881 (CI: 0.869-0.894)** | **0.669 (CI: 0.665-0.673)** | **<0.010** |
| Meropenem | **0.869 (CI: 0.851-0.888)** | **0.711 (CI: 0.705-0.716)** | **<0.001** |
| Ceftazidime | 0.869 (CI: 0.84-0.899) | 0.872 (CI: 0.868-0.875) | 0.849 |
| Piperacillin/tazobactam | **0.873 (CI: 0.858-0.888)** | **0.843 (CI: 0.841-0.845)** | **<0.001** |
| Ceftolozane/tazobactam | **0.968 (CI: 0.958-0.979)** | **0.88 (CI: 0.877-0.883)** | **<0.001** |
| Ciprofloxacin | 0.943 (CI: 0.933-0.953) | 0.950 (CI: 0.948-0.952) | 0.464 |
| Amikacin | **0.962 (CI: 0.955-0.97)** | **0.884 (CI: 0.882-0.886)** | **<0.010** |
| Tobramycin | **0.977 (CI: 0.972-0.982)** | **0.822 (CI: 0.818-0.825)** | **<0.001** |

**Supp. Table 7.** **AUROC comparison between decoupled AMR-GNN and unitig-only model for hold-out test set.** AUROCs are reported as mean with 95% CI.

| **Antimicrobial** | **AMR-GNN** | **Unitig** | ***P*** |
| --- | --- | --- | --- |
| Aztreonam | 0.669 (CI: 0.665-0.673) | 0.667 (CI: 0.648-0.685) | 0.569 |
| Meropenem | **0.711(CI: 0.705-0.716)** | **0.792 (CI: 0.777-0.807)** | **<0.001** |
| Ceftazidime | 0.872 (CI: 0.868-0.875) | 0.858 (CI: 0.843-0.873) | 0.029 |
| Piperacillin/tazobactam | **0.843 (CI: 0.841-0.845)** | **0.89 (CI: 0.878-0.903)** | **<0.001** |
| Ceftolozane/tazobactam | **0.88 (CI: 0.877-0.883)** | **0.922 (CI: 0.909-0.936)** | **<0.001** |
| Ciprofloxacin | 0.950 (CI: 0.948-0.952) | 0.945 (CI: 0.938-0.951) | 0.208 |
| Amikacin | **0.884 (CI: 0.882-0.886)** | **0.845 (CI: 0.832-0.857)** | **<0.001** |
| Tobramycin | **0.822 (CI: 0.818-0.825)** | **0.723 (CI: 0.702-0.743)** | **<0.001** |

**Supp.Table 8.** **Mutations in key AMR-predictive genes for levofloxacin and tobramycin with significantly higher prevalence in non-susceptible compared to susceptible isolates (Chi-square test).**

| **Antimicrobial** | **Gene** | **Effect** | **Non-susceptible isolates (n; %)** | **Susceptible isolates (n; %)** | ***P*** | **References** |
| --- | --- | --- | --- | --- | --- | --- |
| Levofloxacin | *gyrA* | missense_variant c.248C>T p.Thr83Ile | 398 (67.23%) | 9 (2.26%) | <0.001 | [1, 2] |
|  |  | missense_variant c.259G>A p.Asp87Asn | 72 (12.16%) | 3 (0.75%) | <0.001 | [1, 2] |
|  |  | conservative_inframe_deletion&synonymous_variant c.2733_2739delGTCCGAGinsA p.Ser912_Glu913del | 67 (11.32%) | 13 (3.26%) | <0.001 |  |
|  |  | missense_variant c.2707_2709delTCTinsGCG p.Ser903Ala | 66 (11.15%) | 10 (2.51%) | <0.001 | [3] |
|  |  | missense_variant c.2699_2700delCTinsGC p.Ala900Gly | 66 (11.15%) | 10 (2.51%) | <0.001 | [3] |
|  |  | missense_variant c.2679_2680delCCinsGT p.Asp893Glu | 41 (6.93%) | 2 (0.5%) | <0.001 | [3] |
|  |  | missense_variant c.2578G>A p.Gly860Ser | 65 (10.98%) | 10 (2.51%) | <0.001 | [3] |
|  |  | missense_variant c.2011G>A p.Val671Ile | 66 (11.15%) | 10 (2.51%) | <0.001 | [3] |
|  |  | missense_variant c.259G>T p.Asp87Tyr | 12 (2.03%) | 0 (0.0%) | 0.010 | [2] |
|  | *gyrB* | missense_variant c.1404G>T p.Glu468Asp | 13 (2.2%) | 1 (0.25%) | 0.023 | [4] |
|  |  | missense_variant c.1397C>T p.Ser466Phe | 26 (4.39%) | 3 (0.75%) | 0.002 | [4] |
|  | *parC* | missense_variant c.260C>T p.Ser87Leu | 262 (44.26%) | 4 (1.0%) | <0.001 | [2] |
|  |  | missense_variant c.1759G>A p.Ala587Thr | 36 (6.08%) | 11 (2.76%) | 0.024 | [5] |
|  |  | missense_variant c.260C>G p.Ser87Trp | 28 (4.73%) | 0 (0.0%) | <0.001 | [2] |
|  |  | missense_variant c.271G>A p.Glu91Lys | 8 (1.35%) | 0 (0.0%) | 0.049 | [2] |
| Tobramycin | *fusA1* | missense_variant c.1442C>T p.Ala481Val | 10 (1.62%) | 2 (0.15%) | <0.001 | [6] |
|  |  | missense_variant c.1546G>A p.Gly516Ser | 6 (0.97%) | 0 (0.0%) | 0.001 |  |
|  |  | missense_variant c.1763A>G p.Asp588Gly | 25 (4.04%) | 30 (2.18%) | 0.027 | [7] |
|  |  | missense_variant c.1693A>C p.Lys565Gln | 3 (0.48%) | 0 (0.0%) | 0.05 |  |
|  |  | missense_variant c.1683_1686delAGAGinsGGAC p.Glu562Asp | 4 (0.65%) | 0 (0.0%) | 0.014 |  |
|  |  | missense_variant c.1642G>A p.Val548Ile | 3 (0.48%) | 0 (0.0%) | 0.05 |  |
|  |  | missense_variant c.346A>G p.Thr116Ala | 3 (0.48%) | 0 (0.0%) | 0.05 |  |

**Supp. Table 9. Genes previously associated with AMR in *P. aeruginosa*, along with their corresponding positions relative to the PAO1 reference genome (GenBank: NC_002516.1). The list was curated from references [8] and [9].**

| **Gene** | **Start** | **End** |
| --- | --- | --- |
| *gyrB* | 4275 | 6695 |
| *mexR* | 471306 | 471749 |
| *mexA* | 472024 | 473175 |
| *mexB* | 473191 | 476331 |
| *oprM* | 476333 | 477790 |
| *creB* | 523254 | 523943 |
| *creC* | 523943 | 525367 |
| *ampDh3* | 884799 | 885566 |
| *oprD* | 1043983 | 1045314 |
| *oprH* | 1277006 | 1277608 |
| *phoP* | 1277688 | 1278365 |
| *phoQ* | 1278362 | 1279708 |
| *oprF* | 1921174 | 1922226 |
| *parS* | 1950439 | 1951725 |
| *parR* | 1951726 | 1952433 |
| *mexY* | 2208169 | 2211306 |
| *mexX* | 2211322 | 2212512 |
| *mexZ* | 2212677 | 2213309 |
| *galU* | 2215102 | 2215941 |
| *mexS* | 2806350 | 2807369 |
| *mexT* | 2807469 | 2808512 |
| *mexE* | 2808743 | 2809987 |
| *mexF* | 2810009 | 2813197 |
| *oprN* | 2813194 | 2814612 |
| *cprR* | 3450838 | 3451509 |
| *cprS* | 3451506 | 3452801 |
| *gyrA* | 3556427 | 3559198 |
| *arnB* | 3979860 | 3981008 |
| *arnC* | 3981005 | 3982024 |
| *arnA* | 3982021 | 3984009 |
| *arnD* | 3984006 | 3984893 |
| *arnT* | 3984890 | 3986539 |
| *arnE* | 3986536 | 3986883 |
| *arnF* | 3986880 | 3987293 |
| *nalD* | 4006510 | 4007148 |
| *nalC* | 4166518 | 4167159 |
| *dacC* | 4478979 | 4480139 |
| *pbpA* | 4483396 | 4485336 |
| *mpl* | 4498488 | 4499843 |
| *ampR* | 4592990 | 4593880 |
| *ampC* | 4594029 | 4595222 |
| *mexH* | 4706410 | 4707522 |
| *fusA1* | 4769035 | 4771155 |
| *mexV* | 4903466 | 4904596 |
| *mexW* | 4904647 | 4907703 |
| *ftsI* | 4952604 | 4954343 |
| *ampD* | 5064774 | 5065340 |
| *oprJ* | 5149633 | 5151072 |
| *mexD* | 5151078 | 5154209 |
| *mexC* | 5154237 | 5155400 |
| *nfxB* | 5155561 | 5156124 |
| *pmrA* | 5364071 | 5364736 |
| *pmrB* | 5364760 | 5366193 |
| *parC* | 5572222 | 5574486 |
| *parE* | 5576028 | 5577917 |
| *armZ* | 6159560 | 6160699 |
| *ampDh2* | 6176516 | 6177295 |

**Supp. Table 10. Genes previously associated with AMR in Escherichia coli, along with their corresponding positions relative to the E. coli str. K-12 substr. MG1655 reference genome (GenBank: U00096.3). The list was extracted from the Comprehensive Antibiotic Resistance Database (CARD) [10].**

| **Antibiotic Resistance Ontology (ARO)** | **Gene** | **Start** | **End** |
| --- | --- | --- | --- |
| ARO:3004043 | *acrA* | 484426 | 485619 |
| ARO:3003807 | *acrR* | 485761 | 486408 |
| ARO:3003807 | *acrB* | 481254 | 484403 |
| ARO:3003807 | *tolC* | 3178115 | 3179596 |
| ARO:3003378 | *marR* | 1619120 | 1619554 |
| ARO:3004290 | *ampC* | 4377811 | 4378944 |
| ARO:3004055 | *cpxR* | 4104972 | 4105670 |
| ARO:3003900 | *cyaA* | 3991153 | 3993699 |
| ARO:3003370 | *tufA* | 3470145 | 3471329 |
| ARO:3004039 | *emrE* | 568315 | 568647 |
| ARO:3004049 | *fabG* | 1150670 | 1151404 |
| ARO:3004045 | *fabI* | 1350251 | 1351039 |
| ARO:3003386 | *foIP* | 3324063 | 3324911 |
| ARO:3003889 | *glpT* | 2351016 | 2352374 |
| ARO:3003294 | *gyrA* | 2336793 | 2339420 |
| ARO:3003303 | *gyrB* | 3877705 | 3880119 |
| ARO:3004126 | *lamB* | 4247971 | 4249311 |
| ARO:3001328 | *mdfA* | 883673 | 884905 |
| ARO:3004127 | *mipA* | 1865726 | 1866472 |
| ARO:3003775 | *murA* | 3335235 | 3336494 |
| ARO:3003751 | *nfsA* | 891184 | 891906 |
| ARO:3003756 | *nfsB* | 604771 | 605424 |
| ARO:3003390 | *ompF* | 985894 | 986982 |
| ARO:3003308 | *parC* | 3163715 | 3165973 |
| ARO:3003316 | *parE* | 3173504 | 3175396 |
| ARO:3007423 | *ftsI* | 91413 | 93179 |
| ARO:3003899 | *ptsI* | 2534066 | 2535793 |
| ARO:3004109 | *rob* | 4634441 | 4635310 |
| ARO:3003288 | *rpoB* | 4181245 | 4185273 |
| ARO:3003381 | *soxR* | 4277469 | 4277933 |
| ARO:3003511 | *soxS* | 4277060 | 4277383 |
| ARO:3003893 | *uhpA* | 3850136 | 3850726 |
| ARO:3003890 | *uhpT* | 3845776 | 3847167 |

**Supp. Table 11. Genes previously associated with AMR in *Klebsiella pneumoniae*, along with their corresponding positions relative to the *K. pneumoniae* subsp. pneumoniae MGH 78578 strain ATCC reference genome (GenBank: CP000647.1). The list was extracted from CARD [10].**

| **ARO** | **Gene** | **Start** | **End** |
| --- | --- | --- | --- |
| ARO:3003585 | *phoP* | 1292135 | 1292806 |
| ARO:3004041 | *acrA* | 487929 | 489122 |
| ARO:3003373 | *acrR* | 489265 | 489915 |
| ARO:3004580 | *kpnE* | 1743439 | 1743801 |
| ARO:3004583 | *kpnF* | 1743788 | 1744117 |
| ARO:3004588 | *kpnG* | 3312172 | 3313344 |
| ARO:3004598 | *kpnH* | 3313360 | 3314898 |
| ARO:3004598 | *tolC* | 3772816 | 3774294 |
| ARO:3007420 | *lamB* | 4842153 | 4843442 |
| ARO:3007203 | *phoQ* | 1290669 | 1292135 |
| ARO:3003966 | *ompK35* | 1080309 | 1081235 |
| ARO:3003968 | *ompK36* | 2894808 | 2895911 |
| ARO:3004122 | *ompK37* | 1602463 | 1603617 |
| ARO:3007421 | *ftsI* | 103152 | 104918 |
| ARO:3003380 | *ramR* | 622040 | 622621 |

**Supp. Table 12. Genes previously associated with AMR in *Enterococcus faecium*, along with their corresponding positions relative to the *E. faecium* strain SRR24 chromosome reference genome (GenBank: CP038996.1). The list was extracted from CARD [10].**

| **ARO** | **Gene** | **Start** | **End** |
| --- | --- | --- | --- |
| ARO:3003092 | *cls* | 999827 | 1001278 |
| ARO:3003438 | *tuf* | 72500 | 73687 |
| ARO:3003790 | *liaF* | 851722 | 852453 |
| ARO:3003078 | *liaR* | 853524 | 854156 |
| ARO:3003079 | *liaS* | 852450 | 853517 |

**Supp. Table 13. Genes previously associated with AMR in *Staphylococcus aureus*, along with their corresponding positions relative to the *S. aureus* subsp. aureus NCTC 8325reference genome (GenBank: CP000253.1). The list was extracted from CARD [10].**

| **ARO** | **Gene** | **Start** | **End** |
| --- | --- | --- | --- |
| ARO:3003803 | *agrA* | 2096006 | 2096632 |
| ARO:3003074 | *cls* | 2155376 | 2156860 |
| ARO:3003735 | *fusA* | 531032 | 533113 |
| ARO:3003737 | *fusE* | 2310505 | 2311041 |
| ARO:3004661 | *fosB* | 2399531 | 2399950 |
| ARO:3003901 | *glpT* | 330823 | 331263 |
| ARO:3003296 | *gyrA* | 7005 | 9668 |
| ARO:3003301 | *gyrB* | 5034 | 6968 |
| ARO:3003729 | *ileS* | 1107130 | 1109883 |
| ARO:3004572 | *lmrS* | 2246027 | 2247469 |
| ARO:3003917 | *menA* | 951802 | 952740 |
| ARO:3003769 | *mprF* | 1301484 | 1304006 |
| ARO:3003776 | *murA* | 2165268 | 2166533 |
| ARO:3004667 | *norA* | 687425 | 688591 |
| ARO:3003312 | *parC* | 1294197 | 1296599 |
| ARO:3003314 | *parE* | 1292206 | 1294197 |
| ARO:3003323 | *pgsA* | 1214102 | 1214680 |
| ARO:3003285 | *rpoB* | 522301 | 525711 |
| ARO:3003291 | *rpoC* | 525875 | 529471 |
| ARO:3003902 | *uhpT* | 200310 | 201689 |
| ARO:3003794 | *walK* | 25645 | 27471 |
